# Dissecting aneuploidy phenotypes by constructing Sc2.0 chromosome VII and SCRaMbLEing synthetic disomic yeast

**DOI:** 10.1101/2022.09.01.506252

**Authors:** Yue Shen, Feng Gao, Yun Wang, Yuerong Wang, Ju Zheng, Jianhui Gong, Jintao Zhang, Zhouqing Luo, Daniel Schindler, Yang Deng, Weichao Ding, Tao Lin, Reem Swidah, Hongcui Zhao, Shuangying Jiang, Cheng Zeng, Shihong Chen, Tai Chen, Yong Wang, Yisha Luo, Leslie Mitchell, Joel S. Bader, Guojie Zhang, Xia Shen, Jian Wang, Xian Fu, Junbiao Dai, Jef D. Boeke, Huanming Yang, Xun Xu, Yizhi Cai

**Affiliations:** BGI Research-Shenzhen, BGI, Shenzhen, 518083, China; BGI Research-Changzhou, BGI, Changzhou, 213000, China; Guangdong Provincial Key Laboratory of Genome Read and Write, BGI-Shenzhen, Shenzhen, 518120, China; College of Life Sciences, University of Chinese Academy of Sciences, Beijing, 100049, China; Guangdong Provincial Key Laboratory of Synthetic Genomics, Shenzhen Key Laboratory of Synthetic Genomics, Center for Synthetic Genomics, Shenzhen Institute of Synthetic Biology, Shenzhen Institutes of Advanced Technology, Chinese Academy of Sciences, Shenzhen, 518055, China; College of Life Sciences and Oceanography, Shenzhen University, Shenzhen, 518055, China; State Key Laboratory of Cellular Stress Biology, School of Life Sciences, Faculty of Medicine and Life Sciences, Xiamen University, Xiamen 361102, China; Manchester Institute of Biotechnology, University of Manchester, 131 Princess Street, Manchester, M1 7DN, UK; Max Planck Institute for Terrestrial Microbiology, Karl-von-Frisch-Str. 10, 35043 Marburg, Germany; Institute for Systems Genetics and Department of Biochemistry and Molecular Pharmacology, NYU Langone Health, New York, NY 10016, USA; Department of Biomedical Engineering, Johns Hopkins University, Baltimore, Maryland, USA; University of Copenhagen, Universitetsparken 15, 2100 Copenhagen, Denmark; Greater Bay Area Institute of Precision Medicine (Guangzhou), Fudan University, Guangzhou, China; Center for Global Health Research, Usher Institute, University of Edinburgh, Edinburgh, UK; Department of Biomedical Engineering, NYU Tandon School of Engineering, Brooklyn NY 11201

## Abstract

Aneuploidy compromises genomic stability, often leading to embryo inviability, and is frequently associated with tumorigenesis and aging. Different aneuploid chromosome stoichiometries lead to distinct transcriptomic and phenotypic changes, making it helpful to study aneuploidy in tightly controlled genetic backgrounds. By deploying the engineered SCRaMbLE system to the newly synthesized Sc2.0 megabase chromosome VII (*synVII*), we constructed a synthetic disomic yeast and screened hundreds of SCRaMbLEd derivatives with diverse chromosomal rearrangements. Phenotypic characterization and multi-omics analysis revealed that fitness defects associated with aneuploidy could be restored by i) removing most of the chromosome content, or ii) modifying specific regions in the duplicated chromosome. These findings indicate that both chromosome copy number and chromosomal regions contribute to the aneuploidy-related phenotypes, and the synthetic yeast resource opens new paradigms in studying aneuploidy.

**In brief:** Use of SCRaMbLE and newly synthesized Mb-scale Sc2.0 chromosome VII enables insights into genotype/phenotype relationships associated with aneuploidy

**Highlights:**

- *De novo* design and synthesis of a Mb-scale synthetic yeast chromosome VII, carrying 11.8% sequence modifications and representing nearly 10% of the yeast genome.
- A disomic yeast (n + *synVII*) is constructed for dissecting the aneuploidy phenotype
- SCRaMbLE enables systematic exploration of regions causing aneuploidy phenotypes
- Chromosomal copy number and content both contribute to aneuploidy phenotypes
- A 20 Kb deletion on the right arm of synVII leads to fitness improvement linked to up-regulation of protein synthesis

## Introduction

Aneuploidy represents an imbalanced genomic state, in which the copy number of intact or partial chromosomes is altered. At the organismal level, aneuploidy in humans is often intrinsically linked to embryonic lethality particularly in early development with major developmental abnormalities, devastating genetic disorders, tumorigenesis and aging (Baker et al., 2013; Ben-David and Amon, 2020; Holland and Cleveland, 2009; Nagaoka et al., 2012; Pellman, 2007). Therefore, systematic investigation of the underlying molecular mechanisms of aneuploidy is essential to unravel its effects on basic cellular and developmental functions as well as its clinical relevance as a prognostic marker or potential therapeutic target.

Early studies of aneuploid yeast, mice and human cells have unveiled a number of phenotypes and distinct gene expression patterns. Different possible mechanisms have been proposed, attributing the aneuploid phenotypes to changes in many or a small number of critical genes (“mass action of genes” or “few critical genes” hypotheses) (Bonney et al., 2015), resulting in stoichiometric imbalances between different subunits of cellular protein complexes (Chen et al., 2019; Oromendia et al., 2012; Terhorst et al., 2020; Torres et al., 2010a; Tsai et al., 2019).

Although substantial efforts have been devoted to elucidating the causes and consequences of aneuploidy in recent years, the molecular mechanisms underlying the diverse aneuploid phenotypes remain poorly understood. The difficulty in identifying the link between aneuploidy and its associated distinct phenotypes mainly derives from two reasons: 1) the consequences of aneuploidy vary significantly in the context of distinct karyotypes and cell types; 2) limitations of current methods to generate isogenic and stable aneuploid cell populations in multi-cellular organisms (Ben-David and Amon, 2020). Thus, as a unicellular eukaryote, the budding yeast *Saccharomyces cerevisiae* is widely adopted as a simple and suitable model for studying aneuploidy (Mulla et al., 2014). A method has been established to generate aneuploid yeast strains with defined karyotypes by induction of mis-segregation of target chromosomes during mitosis (Beach et al., 2017). However, this method only allows the investigation of immediate consequences of karyotypic changes but fails to further identify the effects of specific regions within a chromosome.

In recent years, DNA synthesis and editing technologies have rapidly evolved and propelled synthetic genomics to center stage (Schindler et al., 2018), best exemplified by the Sc2.0 project that generated a series of synthetic yeast strains bearing designer chromosomes synthesized from scratch (Annaluru et al., 2014; Mitchell et al., 2017; Shen et al., 2017; Wu et al., 2017; Xie et al., 2017; Zhang et al., 2017). As a unique feature of the Sc2.0 genome, the SCRaMbLE (Synthetic chromosome rearrangement and modification by loxP-mediated evolution) system allows the generation of combinatorial genomic diversity through massive rearrangements between designed recombinase recognition sites. This capability has been harnessed for applications including strain improvements for product yield and specific stress tolerance (Blount et al., 2018; Jia et al., 2018; Liu et al., 2018a; Luo et al., 2018; Shen et al., 2018, 2016; Zhao et al., 2020). This on-demand genome rearrangement feature also makes Sc2.0 synthetic yeast a superior model for dissecting the complexity underlying cellular aneuploidy. By generating isogenic aneuploid yeast strains bearing defined Sc2.0 designer chromosomes, SCRaMbLE will enable generating diverse chromosomal rearrangements specifically on the synthetic chromosome(s). Combined genomic, transcriptomic, proteomic and karyotype analyses of the resultant aneuploid yeast strains will shed new light on genotype-to-phenotype relationships associated with aneuploidy.

In this study, we construct the Sc2.0 chromosome VII (*synVII*) and its corresponding disomic yeast to demonstrate the feasibility of this approach. *SynVII* was chosen for two main reasons: first, as the cost of aneuploidy is reported to be proportional to the chromosome length (Tang and Amon, 2013), *synVII* is one of the largest chromosomes at over one million base pairs, representing nearly 10% of the whole yeast genome. Second, few studies conducted in-depth analysis of the cause and consequence of disomic yeast bearing an extra copy of chromosome VII (Torres et al., 2010b). Therefore, our study could facilitate the investigation of multiple aspects of aneuploidy using the disomic *synVII* yeast as a model system. We identify 219 SCRaMbLEd disomic yeasts with massive chromosomal rearrangements specifically limited to *synVII*. Phenotypic characterization and multi-omics analyses reveal two distinct approaches adopted by aneuploid yeast to restore cellular fitness. The substantial fitness cost as the result of aneuploidy can be restored by removing the majority of content from the additional chromosome copy. Interestingly, we found the deletion of a 20 Kb region on the right arm of *synVII* is associated with up-regulation of translation and leads to fitness improvement in varying conditions. Our results indicate that both chromosomal copy number and specific gene content contribute to the aneuploidy phenotypes.

## Results

### The debugged synVII strain exhibits high fitness comparable to the wild-type strain

To utilize SCRaMbLE of synthetic disomic yeast for dissecting aneuploidy phenotypes, we started with the construction of synthetic chromosome VII. *SynVII* was designed following the previously reported Sc2.0 design principles (Richardson et al., 2017), resulting in a final 1,028,952-base pair (bp) synthetic chromosome carrying ∼11.89% modified sequence in comparison with the native sequence.

Three unexpected design features causing significant fitness defects were identified and corrected during the construction process. The synVIIS intermediate strain revealed a significant growth defect when compared to the parental strain synVIIT (as the chromosome was constructed from “right” to “left”) and wild-type (WT) strain BY4741 under optimal growth condition (YPD, 30 °C; **Figure 1A**). By mating the synVIIS strain (*MAT*α) with all 25 single gene knock-out strains (Winzeler et al., 1999) corresponding to the megachunk S region we found that the synthetic *NSR1* gene led to the observed fitness defect (**Figure 1B**). Notably, transcriptome profiling of the synVIIS strain revealed a ∼six-fold up-regulation of *NSR1* transcription compared to synVIIT, which was further confirmed by RT-qPCR analysis, whereas, paradoxically, Nsr1 protein abundance was drastically reduced (**Figures 1C and 1D**). There are several design features of the *NSR1* gene: two synthetic PCRTags within the coding region and one loxPsym site at the 3’UTR region were introduced into synthetic *NSR1*; in addition, the *YGR160W* “dubious ORF” overlapped the *NSR1* gene on its complementary strand, resulting in the insertion of an additional loxPsym site immediately downstream of its stop codon, as dictated by the “rules” of the Sc2.0 design. This resulted in one loxPsym site within the 5’UTR region of *NSR1* (**Figure 1B**). We hypothesized that this “misplaced” loxPsym site might be responsible for the fitness defect. To this end, we individually reverted each designer feature in or near *NSR1* to the wild type counterpart and monitored growth and *NSR1* expression. The steady-state level of *NSR1* returned to the wild-type level by simply removing the loxPsym site at the 5’-UTR region of *NSR1* (**Figure 1E**). In comparison, the removal of the synthetic PCRTags had little effect. These results demonstrate that the loxPsym site at the 5’UTR region of the *NSR1* gene leads to the fitness defect, presumably by interfering with translation since the loxP sequence can form a stem-loop structure (Oliveira et al., 1993). This interpretation accounts for the paradox described previously, namely increased *NSR1* mRNA abundance (presumably a consequence of the reduced protein expression level). Consistent with this finding, the introduction of a loxPsym sequence at the 5’UTR region of *NSR1* resulted in dramatic decreases in protein level and obvious growth defect in the BY4741 strain (**Figure 1E**). A similar pattern was also observed in a synX bug. One loxPsym site was “accidentally” transcribed as a part of *SWI3* 5’ UTR, which led to an increased transcript, but reduced protein level (Zhao et al., 2022). In addition, we noticed that the removal of loxPsym site at the 3’ UTR region could alleviate the cellular defect in the wild-type strain to some extent, suggesting the formation of a potential stem-loop by the two loxPsym sites could further affect the translation. Taken together, significant fitness defect observed in the intermediate synVIIS strain was derived from the loxPsym site inserted into the UTR region of *NSR1*. Another two fitness defects caused by the loxPsym site in megachunk W (**Figure S1A**) and the removal of the tRNA *tN(GUU)G* gene (**Figure S1B**) were identified and corrected, followed by verification of whole genome sequencing, phenotypic, transcriptome and proteome profiling in comparison to the parental strain BY4741. Our results demonstrate that the final version of the synthetic chromosome has minimal effect on yeast cell physiology, and transcriptional and translational states (**Figures S2, 1F and 1G**). Therefore, the synVII strain with its in-depth characterized phenotype is ideal for further aneuploidy studies.

**Figure 1.**
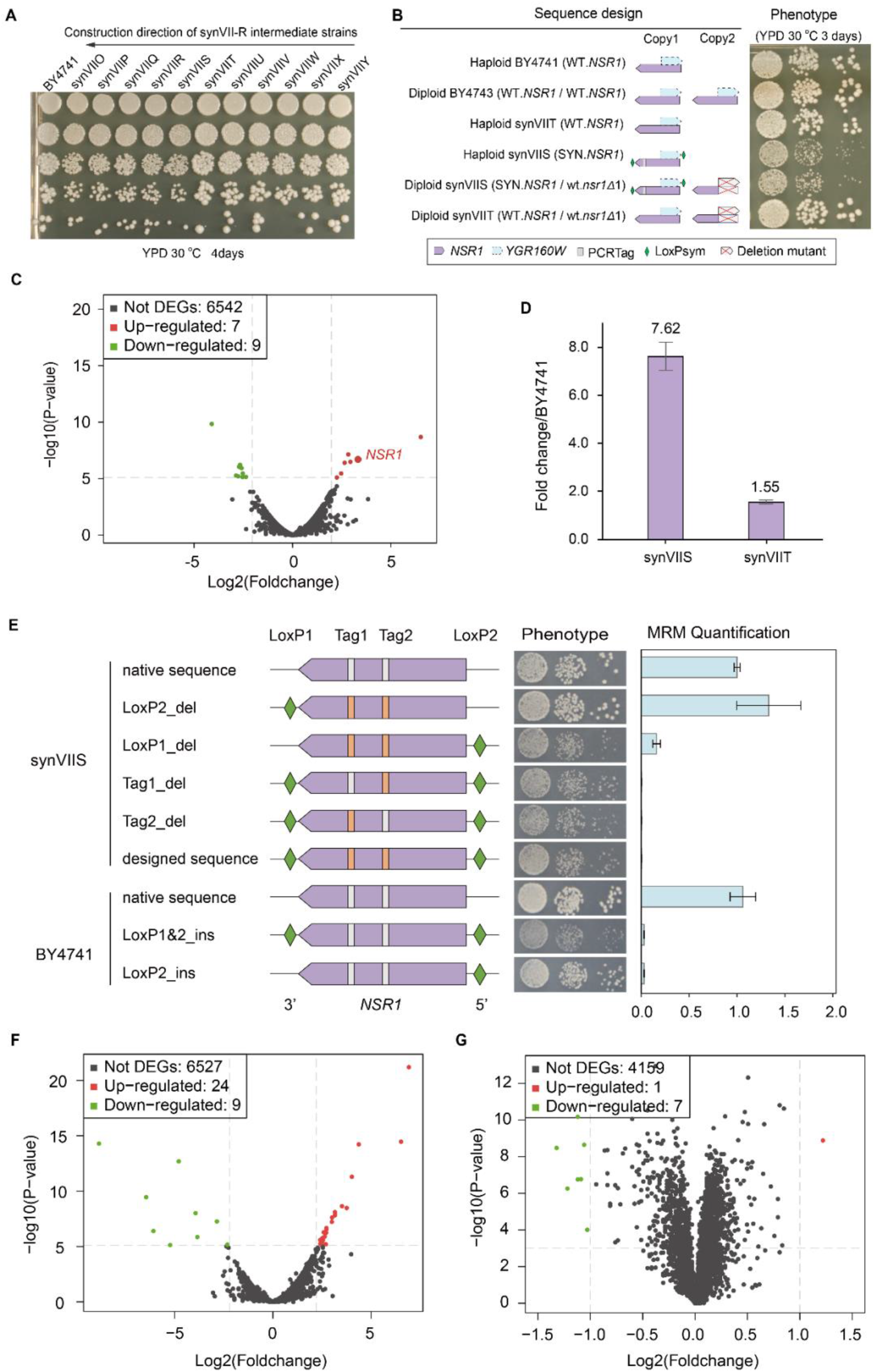
Functional dissection and repair of SynVII. (**A**) Spot assay of synVII-R intermediate strains. The synVII-R is constructed in orientation from megachunk Y towards megachunk O. (**B**) Bug mapping by mating growth-defective query strain synVIIS and its parental strain synVIIT with yeast single knock-out *nsr1Δ1* strain. (**C**) Transcriptome profiling of synVIIS compared to synVIIT reveals significant upregulation of *NSR1* mRNA. Up-regulated features are labelled in red, and down-regulated features are labelled in green. (**D**) qPCR validation of *NSR1* mRNA expression in synVIIS and synVIIT strains. Error bars represent ±SD from three independent experiments. (**E**) Introduction of loxPsym (green) at the 5’UTR region of NSR1 led to a growth defect in both synVII intermediate strain synVII-S and wild type strain BY4741. The corresponding phenotype by plating and protein expression level, quantified by Multiple Reaction Monitoring Mass Spectrometry (MRM-MS) analysis are shown for each constructed strain. Error bars indicate ± SD (n = 3). The white and orange blocks represent wild-type and synthetic PCRTags respectively. Identified dysregulated genetic features at (**F**) transcriptome level and (**G**) proteome level of repaired synVII cells compared to BY4741. Total number of differentially expressed (P < 0.001) features in transcriptome and proteome are presented.

### The *synVII* chromosome in disomic yeast is well maintained and leads to aneuploidy-specific phenotypes

To build a disomic yeast strain with a defined chromosome gain, we took advantage of a well-established system to generate a *chrVII*-specific aneuploid yeast strain (Beach et al., 2017; Hill and Bloom, 1987). The galactose inducible/glucose repressible *GAL1* promoter was inserted adjacent to *CEN7* sequence of WT strain BY4742 to allow controlled inactivation of centromere function, which led to transient nondisjunction of chromosome VII and the generation of the yeast strain YSy140 with two copies of native chromosome VII. YSy140 (*MAT*α) and synVII strain YSy105 (*MAT***a**) were mated and sporulated to obtain the disomic yeast YSy142 bearing one synthetic and one native copy of chromosome VII (**Figure 2A**), which was verified by the whole genome sequencing.

**Figure 2.**
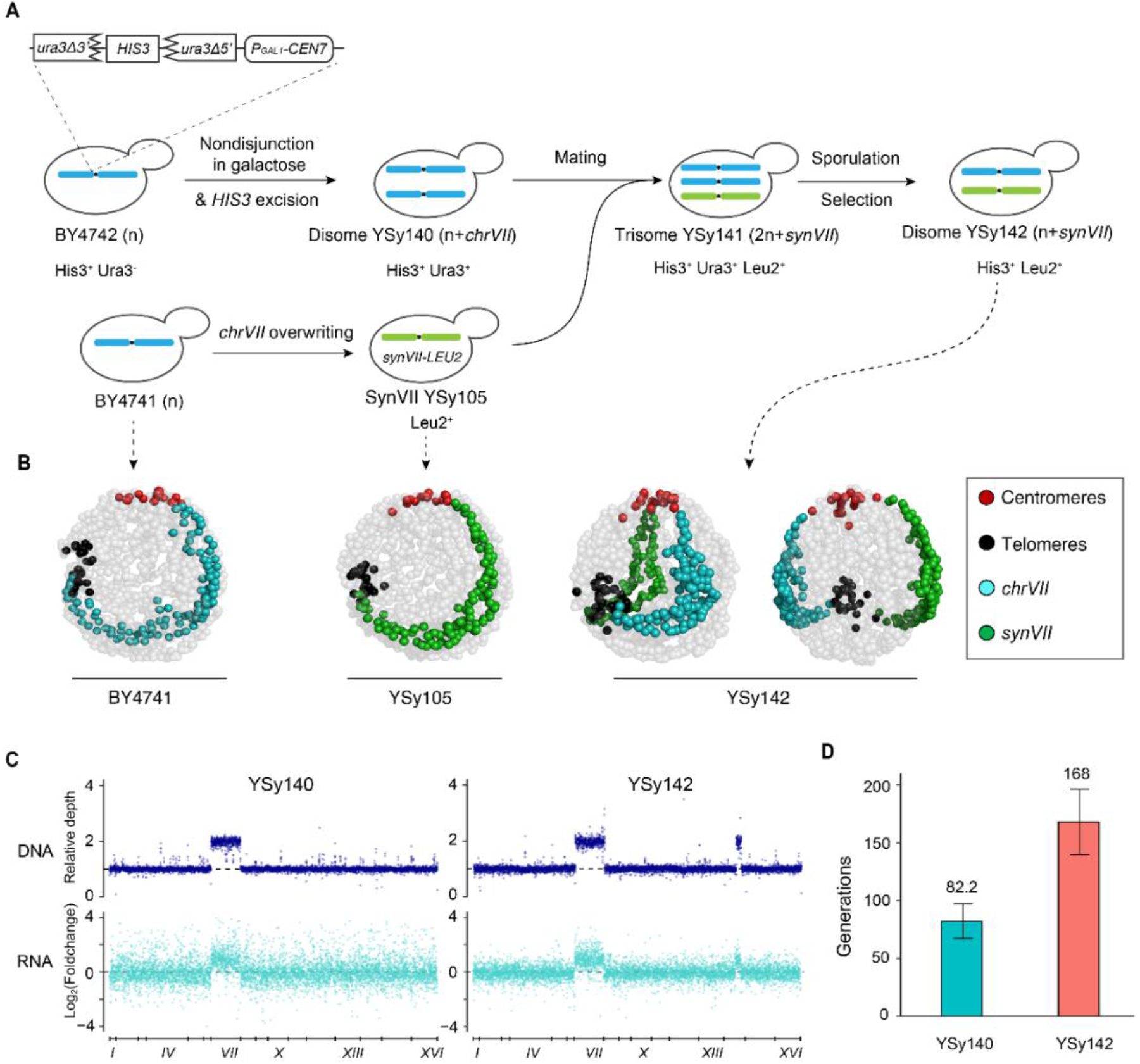
Construction and physiologic analysis of disomic yeasts. (**A**) Schematic illustration of the construction of disome yeast strains YSy140 and YSy142. (**B**) 3D genome organization of native and synthetic chromosome VII in YSy142 strain in comparison with haploid BY4741 and YSy105. Each bead represents a 10 Kb chromosome segment. Centromeres, telomeres, wild type *chrVII* and *synVII* are indicated with red, black, blue and green beads respectively. Other chromosomes are shown in grey. Two angles of view are shown for YSy142 strain. (**C**) The corresponding DNA and mRNA levels track with gene copy number in both disomic yeast strain YSy140 and YSy142. A 60 kb tandem duplication in chromosome XIV is identified in YSy142 and all derived SCRaMbLEd strains. (**D**) Genome stability analysis of YSy140 and YSy142 through long-term growth assays across a time span of ∼220 generations. The number represents the average number of generations maintaining aneuploidy. Error bars indicate ± SD (n = 3).

Next, we systematically examined the consequences of gaining a synthetic chromosome VII in the disome strain YSy142 through detailed analyses of chromosome organization, transcriptional profile, genome stability and phenotypes. By investigating the trajectories of both synthetic and wild-type chromosome VII of YSy142, we found that the two chromosomes are symmetrically arranged in the nucleus (**Figure 2B**). Transcriptome profiling revealed that the majority of genes present on both copies of chromosome VII in YSy142 and YSy140 strains were transcribed (**Figure 2C**). Previous studies have shown that some aneuploid strains are unstable (Potapova et al., 2013). Here we analyzed the stability of YSy142 and YSy140 through long-term growth assays (∼220 generations). Compared to YSy140, YSy142 showed a significant improvement of genome stability. More than 50% of the population of YSy142 stably maintained the *synVII* chromosome over 160 generations. In comparison, the aneuploid strain YSy140 was stable only up to ∼80 generations (**Figure 2D**). The integrity of target chromosomes in both YSy142 and YSy140 was confirmed by whole genome sequencing (**Figure S3**). Overall, the stability of an extra copy of synthetic chromosome VII in the YSy142 strain provides an unprecedented opportunity to dissect chromosome VII specific aneuploidy-associated molecular and phenotypic changes.

To determine how aneuploidy affects the proliferation and physiology, we further characterized aneuploid *synVII* strains YSy140 and YSy142 under conditions with different type of exogenous stress and identified two aneuploidy-specific phenotypes (**Figure S4**). Specifically, disomic strains exhibited increased sensitivity to cycloheximide (a protein synthesis inhibitor) and hydroxyurea (an inhibitor of ribonucleotide reductase) and methyl methanesulfonate (MMS, a DNA damaging agent). These findings are consistent with traits shared by most aneuploid yeast strains harboring different karyotypes reported in previous studies (Torres et al., 2010a; Tsai et al., 2019)

### High-frequency rearrangements revealed in 219 SCRaMbLEd disomic yeasts with varying degrees of fitness recovery

Previous studies demonstrated that extensive unique genotypes could be generated by SCRaMbLE of synthetic chromosomes within populations of Sc2.0 synthetic strains (Blount et al., 2018; Jia et al., 2018; Liu et al., 2018b; Luo et al., 2018; Shen et al., 2016). With the constructed aneuploid yeast strain harboring a complete *synVII*, we sought to determine whether SCRaMbLE could help screen for aneuploid cells which recovered a wild-type phenotype, offering us an opportunity to identify the genes/target regions that drive the aneuploidy-specific fitness defects. To this end, a daughter-cell specific Cre recombinase expression plasmid, pSCW11-*creEBD* (Dymond et al., 2011), was transformed into YSy142 strain to promote SCRaMbLE. After 24 hours, cultured cells were plated onto selective agar medium containing translation inhibitor cycloheximide to select for SCRaMbLEd aneuploid *synVII* derivatives. The dual auxotrophic selection (*Leu2*^*+*^ *Met*^*+*^) of target chromosomes was utilized to maintain the aneuploidy state and ensure that the phenotype is not due to the simple loss of one chromosome copy. The SCRaMbLEd colonies showing improved fitness in the presence of cycloheximide were analyzed by genome sequencing (**Figure 3A**).

**Figure 3.**
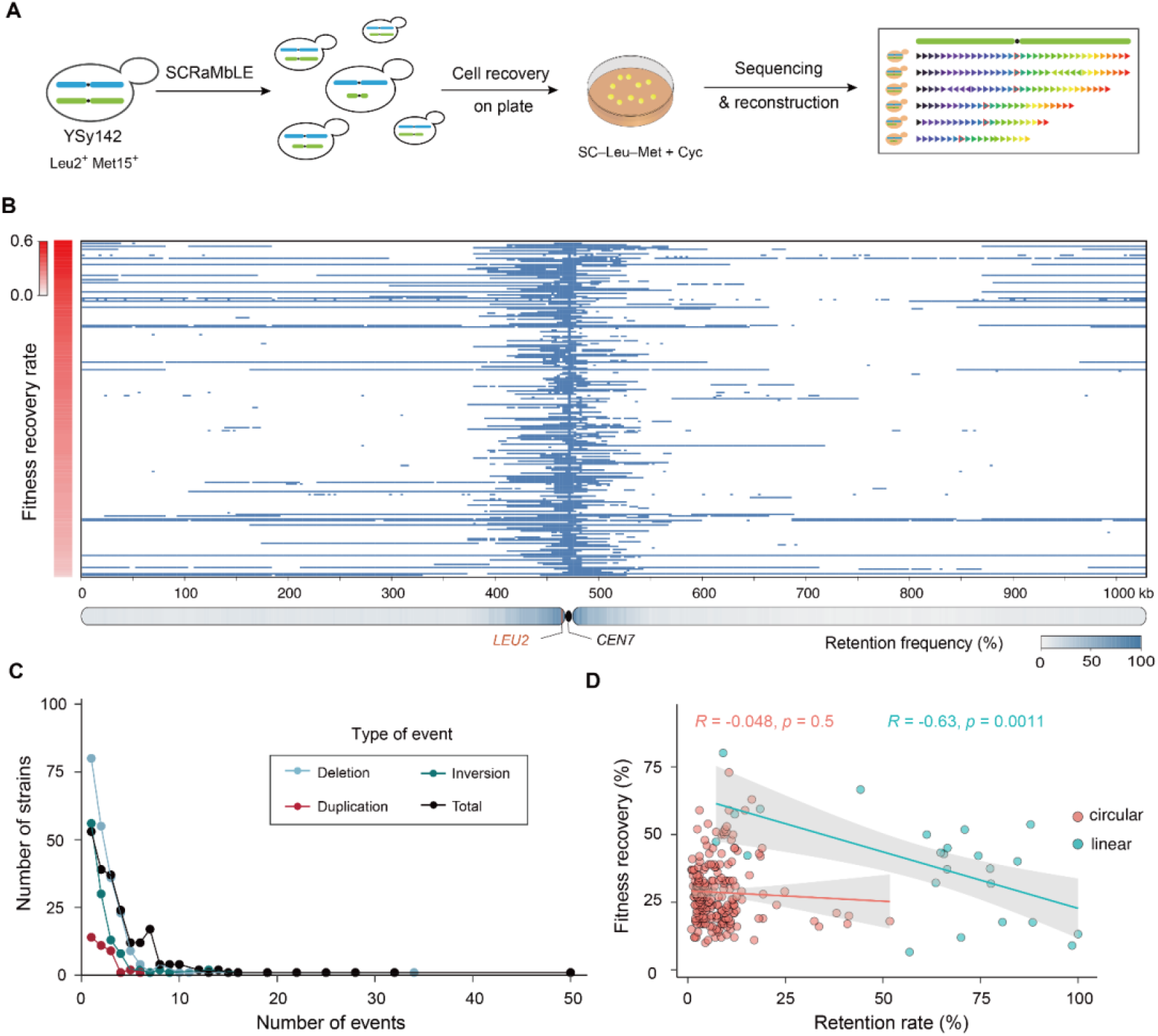
SCRaMbLE of YSy142 disomic yeast. (**A**) SCRaMbLE and analysis workflow. The auxotrophic maker of the YSy142 strain was swapped from *HIS3* to *MET15*. See method “Construction of aneuploid *synVII* strain” section for details. (**B**) The fate of each segment flanked by two loxPsym sites in each strain is indicated as reserved (light blue) or deleted (white) by any SCRaMbLE event. Y-axis shows the relative average phenotypic recovery rate of each SCRaMbLEd strain in comparison to that of YSy142 represented by color scale (n ≥ 200). (**C**) The distribution of recombination events for all selected SCRaMbLEd aneuploid strains. The number of events per strain has a long tail with some strains having 22 and 25 distinct recombination events. (**D**) Correlation between chromosome retention rate and fitness recovery in SCRaMbLEd strains with circular *synVII* and SCRaMbLEd strains with linear synVII. Each dot represents one SCRaMbLEd strain. R indicates the Pearson’s correlation. Solid line: fitted curve (ggplot:geom_point), geom_smooth (method=lm, se=TRUE). Gray area: 95% confidence range for fitted curve.

In all 219 selected SCRaMbLEd strains, the fitness recovery rate was calculated by comparing average colony size (from ≥ 200 single colonies) of each strain to that of the unSCRaMbLEd aneuploid yeast parent YSy142. Our results revealed varying degrees of fitness recovery rate from 6.6% to 80% for SCRaMbLEd strains harboring distinct content and size of *synVII* (**Figure 3B**). Not surprisingly, a peak of sequence retention was found within *CEN7* adjacent regions due to the maintenance of selection for prototrophy (*Leu2*^*+*^). However, from the whole genome sequencing, we observed an unequal proportion of reads of wild-type versus synthetic *synVII* PCRTags for three SCRaMbLEd strains, with the relative ratio at around 2:1 (**Figure S5A**). Presumably, the ratio should be equally distributed since there is one copy of wild-type and synthetic chromosome VII each in the disome yeast. Further analysis by flow cytometry on one of the three strains confirmed that it is a trisomic yeast (**Figure S5B**), possibly resulted from spontaneous whole-genome duplication post or during SCRaMbLE. Thus, these three strains are excluded from the collection of selected SCRaMbLEd strains for further analysis since their phenotypes might also be influenced by the corresponding genome ploidy as suggested in previous studies (Sheltzer et al., 2011).

Among the 969 recombination events that occurred in cells that showed recovery of cellular fitness, three types of events including deletion, inversion and duplication, were observed at the frequencies of 62.2%, 29.2% and 8.6% respectively. A recent SCRaMbLE study suggested that the recombination frequency in a haploid synthetic yeast was not as high as that in the synIXR experiment (Jia et al., 2018). It is very likely that the random recombination between two loxPsym sites in a region containing essential genes in haploid yeast would lead to lethality and consequently resulted in bias of recombination events, although the random nature of SCRaMbLE events can be preserved in heterozygous diploids (Shen et al., 2018). Presumably, in our study we might observe an increase of recombination frequency since SCRaMbLE only occurs in the synthetic chromosome, while the native copy remains intact. As expected, for each SCRaMbLEd strain, we observed an average number of events at 4.42. We also observed a long tail in the distribution of events per strain, with ≥10 events found in 17 strains, with two strains containing 32 and 50 events, respectively (**Figure 3C**).

In general, we observed a negative correlation between chromosome retention rate (the frequency of each *synVII* segment preserved after SCRaMbLE across all selected SCRaMbLEd strains) and fitness recovery rate (the relative colony size of each SCRaMbLEd strain compared to YSy142 under the same culturing condition) (**Figure 3D**). We found 31 SCRaMbLEd strains carrying circular chromosomes with higher fitness recovery rates (40% to 60%) tended to lose most of both *synVII* chromosome arms, retaining only 1% to 19% of the original chromosome arms, namely the centromere adjacent regions. In contrast, 18 out of the 24 strains with >50% of retention rate showed only moderate fitness recovery rates (averaging 32.4%). Our result supports the idea that gene dosage is contributing to aneuploidy and by reducing the copy number via SCRaMbLE the fitness defect of disomic yeast is rescued.

### The frequency of circular *synVII* is significantly higher than that of linear *synVII* maintained in SCRaMbLEd disomic strains

We identified two types of *synVII* chromosome structural conformations (circular and linear forms) in the 219 SCRaMbLEd disomic yeast. Interestingly, the frequency of generating circular SCRaMbLEd synthetic chromosome VII was surprisingly high. Around 89% of all selected SCRaMbLEd disomic strains (in total 195) maintained circular SCRaMbLEd *synVII* with sizes ranging from 10 Kb to 532 Kb, while only 11% of selected strains (in total 24) carried the original linear SCRaMbLEd *synVII* with size ranging from 74 Kb to 1028 Kb. One possible explanation for this is as follows. Once a circularized chromosome forms, which requires a single intramolecular SCRaMbLE event, it is “primed” to give rise to daughter deletions that remove both chromosome arms in a single step. Moreover, whereas linear *synVII* can continually give rise to additional daughter circles, once locked into the circular state it cannot return to a linear state via SCRaMbLE. Another potential explanation is that some (or several) genes near one of the telomeres are very toxic in multiple copies. Thus, the formation of circular chromosomes would remove this gene preferentially and may have been selected for.

We observed that the average coverage depth of circular SCRaMbLEd *synVII* was ∼3 times lower than the native copy (**Figure S6**). By both flow cytometry measurement and sporulation analysis, we have confirmed that the depth difference is not an artifact of derived strains that become diploid trisomes triggered by spontaneous whole-genome duplication (**Figure S7**). It is possible that circular *synVII* is not stable and lost in a subpopulation, raising the concern that the improved fitness is due to the average lowered copy number in the mixed population. To exclude this possibility, we selected three representative samples with distinct circular *synVII* contents from the 195 strains for long-term genome stability assays. Our results showed that the genome of SCRaMbLEd aneuploid strains is fairly stable. More than 50% of the population can stably maintain the circular SCRaMbLEd *synVII* chromosome over 60 to 200 generations in the absence of selection (**Figure S8**). Considering at the time (∼3 days culturing) of phenotypic assays and sampling for genome sequencing, more than 90% of the population in each selected strain still maintained the SCRaMbLEd *synVII* chromosome, we conclude that the apparent low copy number of circular SCRaMbLEd *synVII* is not the main reason for the observed fitness recovery. It has been previously reported that the average read depth for non-synthetic nuclear chromosome is greater than that for the synthetic circular chromosome arm, indicating higher recovery of linear versus circular chromosomes during the sample preparation process for sequencing (Dymond et al., 2011; Shen et al., 2016), potentially explaining the observed lower depth of circular SCRaMbLEd *synVII*.

In contrast to SCRaMbLEd strains bearing circular *synVII*, the recovery rates of screened strains that retained linear *synVII* exhibit a relatively wide range (from 6.6% to 80.2%), with an average recovery rate of around 39.1%. Using a previously established method (Shen et al., 2016), we reconstructed the full SCRaMbLEd *synVII* chromosome sequence of all 24 screened strains (**Figure S9A**). In 22 out of 24 strains with varying recovered phenotype, *synVIIL* was well maintained, showing few SCRaMbLE events had occurred, while the chromosome content of *synVIIR* across these strains changed drastically. The most frequently observed recombination event was deletion, at a frequency of 61.9%, followed by inversion at a frequency of 34.1%. We further explored the deletion distribution across the entire chromosome and observed a deletion hotspot in the right arm region of *synVII* (**Figure S9B**). For the top 18 strains with relatively high fitness improvement (the corresponding average recovery rate >30%), we visualized the copy number of each segment between two adjacent loxPsym sites in the original order of *synVII*. Large fragment loss within the middle half of chromosome right arm and some small deletions also located in the left arm represented the most common deletion types (**Figure S9C**).

### The loss of majority chromosome content by SCRaMbLE leads to significant fitness improvement of disomic yeast with reversed Environmental Stress Response

Strains losing 85% of the chromosome content represent the leading type among all selected 195 SCRaMbLEd strains with circular *synVII*, at a frequency of 91% (in total 178 strains), but the recovery rates of these strains varied significantly. We speculate that the retained chromosome contents include favored gene combinations that restore fitness under specific stress conditions, and thus the fitness of strains with varying retained gene contents would differ. We then selected the top 18 strains with recovery rate above 46% and examined their phenotypes under normal condition (YPD) as well as stress conditions related to DNA replication and repair, translation, and osmolarity regulation. Compared with the parental YSy142 strain, we observed a general improvement of fitness in all 18 strains, but to varying degrees (**Figure 4A**).

**Figure 4.**
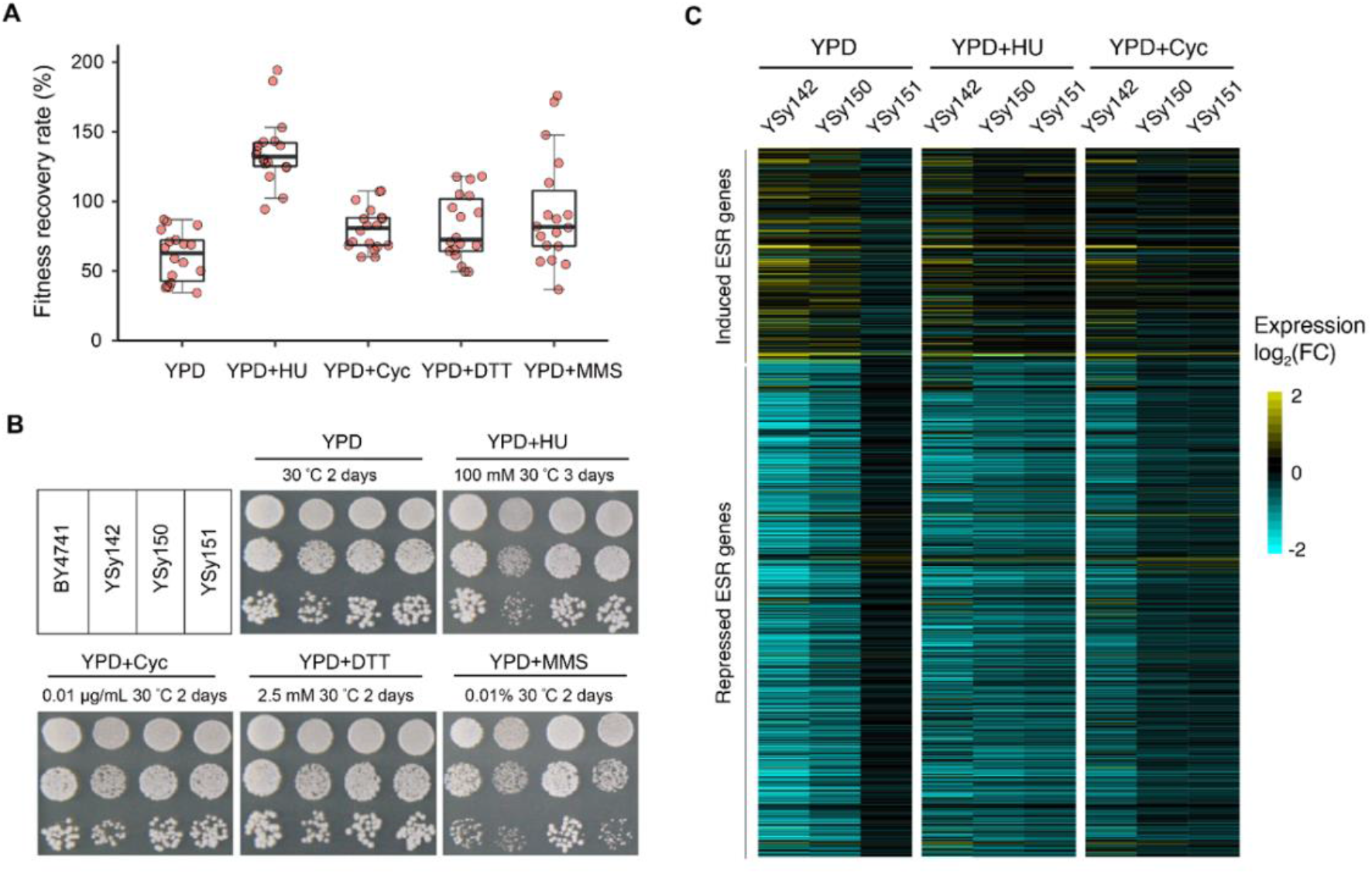
Growth assay and ESR profiling of disomic yeast with circular SCRaMbLEd *synVII*. (**A**) General improvement of fitness at varying degrees was observed for the top 18 SCRaMbLEd strains in five representative conditions. Each dot represents average fitness recovery rate of one SCRaMbLEd strain calculated based on multiple single colonies (n ≥ 200). (**B**) Spot assay of two representative strains of the 18 SCRaMbLEd strains under various conditions, showing general improved fitness. (**C**) The ESR genes expression profile of the unSCRaMbLE strain YSy142 and two SCRaMbLEd strains (YSy151 and YSy150) after normalization with the reference wild type strain BY4741 in three selected conditions. The fold change is represented by the color scale (yellow: up-regulated; blue: down-regulated). Conditions include: YPD at 30 °C for 2 days; YPD + cycloheximide (Cyc, 0.01 μg/mL) at 30 °C for 2 days; YPD + DL-Dithiothreitol (DTT, 2.5 mM pretreatment for 1 h) for 2 days; YPD + hydroxyurea (HU, 100 mM) at 30 °C for 4 days; YPD + methyl methane sulfone (MMS, 0.01% v/v) at 30 °C for 3 days; YPD, yeast extract peptone dextrose.

Previous studies described that exponentially growing disomic yeast strains typically exhibited a gene expression pattern designated as yeast environmental stress response (ESR), which includes the up-regulation of ∼300 genes and the down-regulation of ∼600 genes (also known as the “induced (i)ESR” and “repressed (r)ESR” respectively) (Terhorst et al., 2020; Torres et al., 2007). Our transcriptome analysis consistently revealed that the disomic YSy142 also exhibited the ESR transcriptional signature. To be more specific,>70% of the reported iESR genes are up-regulated and>80% of the rESR genes were down-regulated in YSy142. Because the ESR pattern is highly correlated with the fitness of disomic yeast strains under stressful conditions (Sheltzer et al., 2012), we hypothesized that the SCRaMbLEd strains with recovered fitness would show a reversed ESR pattern. The fitness of both YSy150 and YSy151 SCRaMbLEd strains, which lost 95.2% and 88.4% of synthetic chromosome VII respectively, was significantly improved under most tested conditions compared to YSy142 (**Figure 4B**). Thus, these two strains were chosen for further transcriptome analysis with a focus on ESR-related genes. As expected, our results revealed a reversed trend of the transcriptional signature in both strains under three selected representative conditions, in which most of the iESR/rESR genes no longer showed significant up-/down-regulation in comparison with YSy142 (**Figure 4C**). These results demonstrate that the degree of the ESR correlates well with cell proliferation rate and the SCRaMbLE process is efficient and effective to recover aneuploidy specific fitness. Taken together, our results indicate that aneuploid proliferation defects can be driven by gene changes across the entire chromosome (Bonney et al., 2015).

### The deletion of a 20 Kb region on *synVII* is linked to up-regulation of translation and leads to fitness improvement

In contrast to SCRaMbLEd strains with circular *synVII*, the SCRaMbLEd strains bearing linear *synVII* maintained the majority chromosome content (chromosome retention rate ranging from 7.2% to 100%) and showed a relatively minor fitness recovery rate. We aimed to disclose whether a distinct mechanism other than the “mass action of genes” hypothesis was utilized by these yeast cells to cope with the aneuploidy-induced stress. Thus, we performed comparative proteomics analysis on 5 disomic strains with linear SCRaMbLEd *synVII* that exhibited varying recovery rates grown under normal (YPD) as well as stress conditions (YPD supplemented with cycloheximide and hydroxyurea) (**Figure 5A**). In addition, we observed that protein synthesis and ribosome biogenesis processes were significantly up-regulated in all selected strains under various conditions, suggesting that the increased activity of protein biosynthesis was linked to the improved fitness of disomic strains.

**Figure 5.**
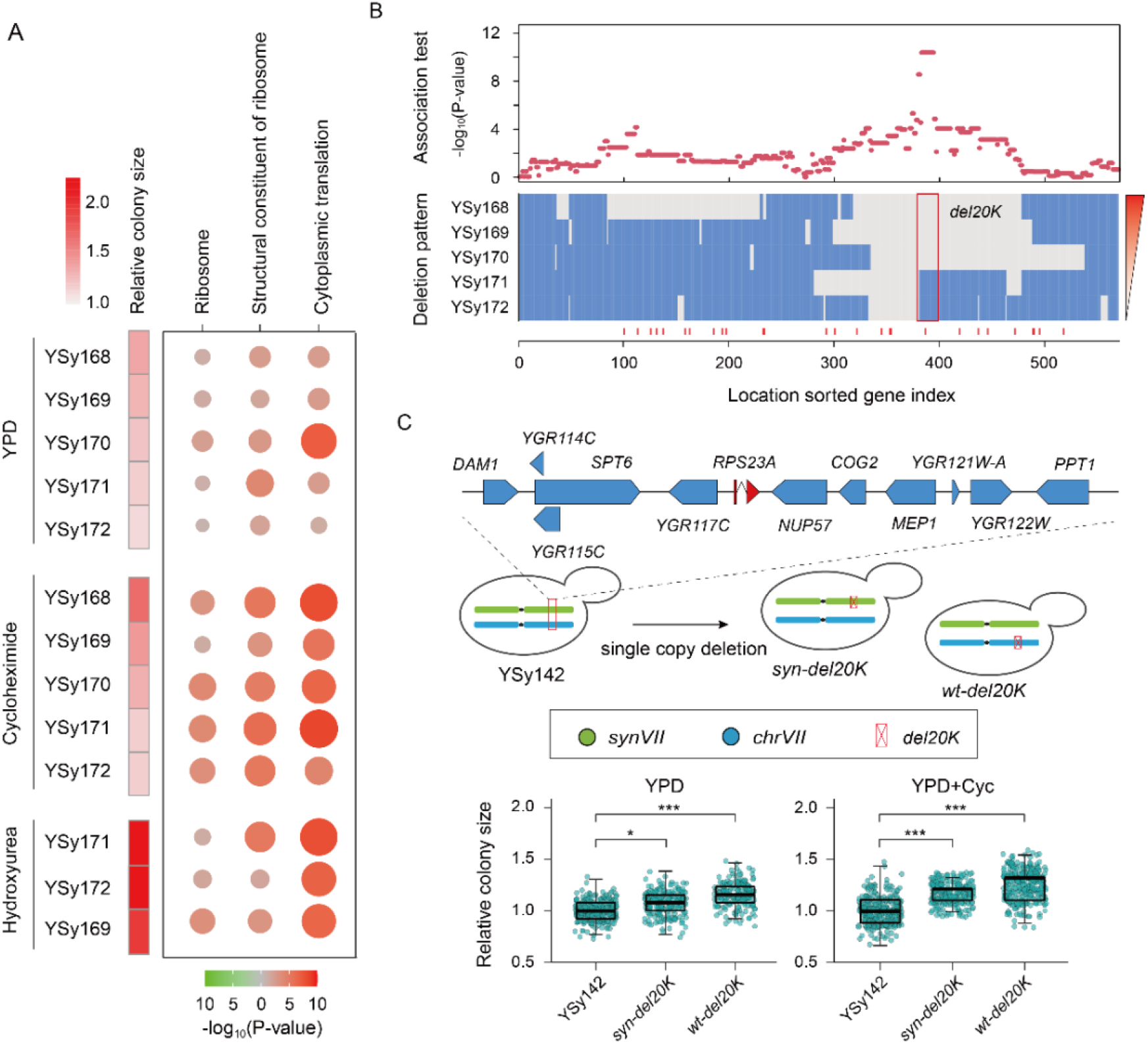
The improvement of *synVII* aneuploidy phenotypes by the deletion of a 20 Kb region (del20K) on *synVIIR* is associated with the upregulation of protein biosynthesis. (**A**) The proteomics analysis of 5 disomic strains with linear SCRaMbLEd *synVII* exhibited varying recovery rates grown on YPD and two stress conditions. (**B**) The chromosome-wide association analysis against the 5 disomic strains reveals a 20 Kb region which might play an essential role in the observed fitness improvement. Each red dot represents the statistical significance of each gene for the association between its deletion pattern and relative colony size. The 5 selected strains are ranked by their corresponding fitness recovery rate. The regions deleted and reserved after SCRaMbLE is presented in grey and blue respectively. (**C**) The phenotypic assay shows that the fitness can be improved by recreating the del20K deletion of *synVII* or wild-type *chrVII* in YSy142. Each dot represents the relative colony size of each colony compared to YSy142 (n ≥ 200). * P < 10E-10, *** P < 10E-30. YPD + cycloheximide (Cyc, 0.01 μg/mL) at 30 °C, YPD + hydroxyurea (HU, 100 mM) at 30 °C.

Next, we wanted to determine the correlation among the observed rearrangement events on linear *synVII*, enhanced expression of the translation machinery and improved phenotype. Based on the observed deletion hotspot on *synVII* and distribution of genes involved in translation and ribosome biogenesis, we looked for the key rearrangements that were potentially related to fitness improvement of disomic yeast strains after SCRaMbLE. We also performed a chromosome-wide association analysis to study the recombination effect of every single gene on cell fitness and found that the deletion of several genes might be involved in fitness improvement (**Figure 5B**). In particular, a ∼20 Kb deletion (del20K) involving 12 genes in *synVII* chromosome showed the strongest association with fitness recovery (P = 4.20E-11). We speculate that genes located in the del20K are likely responsible for the observed recovered phenotype of selected SCRaMbLEd strains (**Figure 5C**). To further verify the causality of the del20K on the fitness phenotype, we re-created the del20K deletion on *synVII* or wild-type chromosomes in YSy142 for further phenotypic analysis. By deleting one copy of either wild-type or synthetic del20K, we observed a general fitness improvement (**Figure 5C**). Particularly, the average cellular fitness recovery rate of the corresponding resulting strain in the presence of cycloheximide is 16.6% (P = 1.05E-50) and 25.4% (P = 1.03E-86) respectively. Our results show that the 20 Kb deletion indeed contributes to the improvement of aneuploidy phenotypes. Although the underlying molecular mechanism is not entirely clear, our result suggests that the chromosome VII genetic effect of yeast colony size exists at different locations of the chromosome in different conditions, so that a systematic up-regulation of the protein synthesis process potentially associated with the del20K could overcome the genetic effect causing *synVII* aneuploidy phenotypes.

## Discussion

Here we describe a novel system for studying the consequences of varying aneuploidy states in a controllable manner by SCRaMbLEing constructed aneuploid Sc2.0 yeast strain. In general, it is well-known that different aneuploid states can lead to distinct patterns of phenotypic and gene expression profiles and aneuploid cells bearing large chromosomal lesions often affecting hundreds of genes (Beach et al., 2017; Bonney et al., 2015; Pavelka et al., 2010; Torres et al., 2007). Following traditional approaches which are incapable of micro-manipulating the target chromosome content, it is extremely challenging, if not impossible, to systematically characterize the consequences and explore the causality of these phenotypes. In our study, we have demonstrated that this technical bottleneck was overcome by the form of the developed inducible aneuploidy system, which enables controlled perturbation(s) of the specific chromosomal region(s) and has two advantages. Except for synthetic chromosome VII, the aneuploid strain YSy142 maintains the same genetic background as its parental haploid strains BY4741 and YSy105, thereby enabling the analysis of phenotypic consequences of aneuploidy in a consistent genetic background. In addition, the unique built-in SCRaMbLE design limits perturbation of the chromosomal structure and content to the synthetic chromosome, and consequently the investigation of the potential causal regions for aneuploid-specific phenotypes in a large chromosome of a defined karyotype is readily achieved.

We demonstrated the feasibility of using the disomic *synVII* strain YSy142 for studying aneuploidy and identified over two hundred SCRaMbLEd aneuploid strains for in-depth analysis. Our result shows that losing the majority content of *synVII* chromosome can relieve the cell from fitness defects under stress conditions, suggesting that “mass action” of genes across the entire chromosome VII is mainly responsible for the proliferation defect associated with aneuploidy. In addition, a 20 Kb deletion hotspot on the middle half of the right arm of linear *synVII* chromosomes of SCRaMbLEd strains was associated with fitness recovery. We further reconstructed this structural variant in disomic yeast strain YSy142 and demonstrated that the del20K suffices as responsible for the observed fitness recovery. Proteomics analysis suggests that the potential underlying mechanism for this is the up-regulation of protein biosynthesis. These findings uncover the potential critical roles of one or more of these genes within del20K that could contribute to aneuploidy-induced stress via interfering with the protein synthesis process. By simply removal of the second copy of these genes on *synVII* chromosome, cell fitness could be alleviated. It will be of interest to dissect this region to further identify its key regulator(s). Interestingly, we also found that the significant fitness recovery for three SCRaMbLEd aneuploid strains is due to spontaneous whole-genome duplication (WGD). By flow cytometry, we confirmed that these three SCRaMbLEd aneuploid strains that present near wild-type fitness recovery have experienced a homozygous increase in the DNA content from disome (n+1) to 2n+1. It is known that yeast cells can undergo spontaneous alterations of cell ploidy to gain a growth advantage under stressful conditions (Harari et al., 2018). Thus, we speculate that the observed spontaneous WGD was a compensatory response to the increased fitness burden triggered by aneuploidy, and furthermore, it suggests that “dialing down” relative expression of the offending genes by half suffices to greatly reduce the fitness defect.

To summarize, the inducible aneuploidy system developed in this study holds great potential to be further applied to systematically construct a full 16-chromosome collection of aneuploid yeast strains heterozygous for synthetic chromosomes and dissect the molecular mechanisms underlying how aneuploidy impacts cell physiology and disease states. In light of the recent completion of all Sc2.0 yeast chromosomes and the prospect of *de novo* assembly of chromosomes from other species including humans, our strategy can potentially be applied to study aneuploidy in diverse chromosomal, cellular and species contexts. With the technical feasibility of *de novo* designing and constructing Mb-scale chromosomes combined with the flexibility of genomic rearrangements conferred by SCRaMbLE, we envision our strategy will revolutionize synthetic genomics and aneuploidy studies.

## Data and code availability

The data that support the findings of this study have been deposited into CNGB Sequence Archive of CNGBdb with accession number CNP0002230.

## Acknowledgments

The project was funded by National Key Research and Development Program of China (No.2018YFA0900100). This work was also supported by UK Biotechnology and Biological Sciences Research Council (BBSRC) grants BB/M005690/1, BB/P02114X/1 and BB/W014483/1, and a Volkswagen Foundation the “Life? Initiative” Grant (Ref. 94 771) to YC. JD is supported by a Royal Society Newton Advanced Fellowship (NAF\R2\180590) hosted by YC. This work was also funded by National Natural Science Foundation of China (31800078 and 21901165), Science, Technology and Innovation Commission of Shenzhen Municipality under grant No. JCYJ20180507183534578, Guangdong Provincial Key Laboratory of Genome Read and Write (No. 2017B030301011), Guangdong Provincial Academician Workstation of BGI Synthetic Genomics (No. 2017B090904014) and Shenzhen Engineering Laboratory for Innovative Molecular Diagnostics (DRC-SZ[2016]884). We thank the DNA assembly automation platform of China National Genebank for the support on synthetic chunk assembly. LAM, JSB and JDB were supported by grants from the US National Science Foundation.

## Author contributions

Conceptualization, Y.S., Y.C.; funding and resources, Y.S., Y.C.; data production, F.G., Y.W., Y.S., J.Z., Z.L., D.S., Y.D., W.D., T.L., R.S., H.Z., S.J., C.Z., S.C., T.C. and Y.W.; data analyses, investigation, and visualization, Y.W., J.Z., J.G., Y.L., L.M., J.B., G.Z., X.S. and X.F.; writing – original draft, Y.S., X.F., Y.W., Y.R., J.Z., F.G., Y.C. and J.B.; writing, review & editing: all co-authors.

## Materials and Methods

### *SynVII* design and construction

Methods of synthetic chromosome design, synthesis and assembly described previously were used in this study (Richardson et al., 2017; Shen et al., 2017). The final version of *synVII* is defined as yeast_chr07_3.57, with a total of ∼11.89% sequence been modified. The sequence of synthetic chromosome *VII* was computationally segmented by BGI customized software “Segman” to 25 ∼50 kb megachunks, then to 129 ∼10 kb chunks and to final 485 ∼3 kb minichunks for synthesis from scratch. More information on *synVII* design can be accessed on the synthetic yeast project website (www.syntheticyeast.org) and **Table S1**. A 2572 bp homologous region with I-*Sce*I site was designed on both semi-synthetic *synVII*s (*synVII-L* and *synVII-R*, strain ID: YSy088, YSy089) for integrating into the full-length *synVII* chromosome. The overlap region shared between adjacent chunks was designed to be ∼800-1200 bp long. For megachunk integration, the ligation step was skipped and all 5-6 chunks (equal to 1 megachunk) were directly co-transformed into yeast to replace the corresponding wild-type sequence using homologous recombination followed by selection. Strains generated in this study are listed in **Table S2 &S3**.

### Stress sensitivity assay

Spot dilution assays were previously described (Shen et al., 2017). Single colonies of BY4741, BY4742, and synVII (yeast_chr07_9.05, strain ID: YSy107) were cultured overnight in YPD medium at 30 °C. Cells were 10-fold serial diluted and spotted onto various selective plates, with YPD medium plates at 30 °C as control. All plates with drugs or adjusted pH (pH 4.0 and pH 9.0) were incubated at 30 °C for 2-4 days. Plates were incubated at 25 °C and 37 °C for temperature stresses.

### Omics analyses

Paired-end whole genome sequencing was performed on the synVII (yeast_chr07_9.05, strain ID: YSy107) with the BGIseq500 platform. A 200-400bp sequencing library was prepared according to BGI’s DNA preparation protocol using the MGIEasyTM Universal DNA Library Prep Kit V1.0 (catalog number: 1000006986). The YSy107 and BY4741 strain both with 3 biological replicates were prepared using sample preparation methods established previously for transcriptome and proteome analysis (Shen et al., 2017). For proteomics, proteins were labeled by iTRAQ reagent-8plex multiplex kit (catalog number: 4381663; BY4741 labeled in isobaric tag: 114, 115, 117, 121; YSy107 labeled in isobaric tag: 118, 119, 121) and the peptides were fractionated with high pH RP method and analyzed by a Q Exactive™ HF-X mass spectrometer (Thermo Fisher Scientific, San Jose, CA) coupled with an online HPLC.

Sequencing reads QC, data processing and analysis were performed as described previously (Shen et al., 2017). For genome sequencing analysis, after filtering low-quality reads with SOAPnuke (Chen et al., 2017), clean reads were mapped to a reference sequence of the *synVII* yeast genome using Bowtie2 2.2.5 with default parameters (Langmead et al., 2009). Both GATK3.8 (McKenna et al., 2010) and SAMtools (Li et al., 2009) pipelines were used to identify the variants, which were further validated by Tablet (Milne et al., 2013). For transcriptomics, reads were mapped to genomes by TopHat (Trapnell et al., 2009), and differentially expressed genes were analyzed by DEseq2 (Anders and Huber, 2010). For proteomics, Mascot and IQuant (Wen et al., 2014) were used for protein identification and quantification. Gene enrichment and coexpression enrichment analyses were performed using KEGG pathways and Gene Ontology annotations.

### Fitness defect mapping and reversion

The well-established “synthetic genetic array” (SGA) analysis method, in which a query mutant is crossed with a pre-designed yeast gene-deletion mutant library (Tong et al., 2001; Winzeler et al., 1999) was applied for defect mapping in synVIIS intermediate strain (strain ID: YSy093). In Megachunk S, 20 of the 23 genes in Megachunk S have corresponding gene-deletions in the ∼5000 yeast gene-deletion mutant SGA library (the remaining 3 are essential genes) and were recovered from the library. These mutant strains were mated with the synVIIS strain as well as a control synVIIT strain. By mating these deletion strains individually with the defective synVIIS strain, the resulting diploid strains with one gene copy deleted and another copy that cannot maintain proper function will pinpoint the cause of the observed growth defect namely *YGR159C*.

For the defect observed in megachunk W, YSy100 bearing full synthetic megachunk W was backcrossed to wild-type strain BY4742, followed by sporulation and tetrad dissection. The generated spores were then selected for further analysis by previously described method “PoPM” (Wu et al., 2017) and this helped to pinpoint the defect to synthetic variant sequences in *MTM1* and *RAD2* genes. Each of the designed sequence variants were individually introduced into the BY4741 strain to further examine their effects.

### Construction of aneuploid *synVII* strain

The construction of aneuploid *synVII* strain was performed based on a previously described method involving transient nondisjunction of the target chromosome (Anders et al., 2009). In this study, a conditional centromere construct (*P*_*GAL1*_*-CEN7*) and a *ura3Δ3’-HIS3-ura3Δ5’* cassette were introduced on chromosome VII in the BY4742 strain (**Figure 2A**). The conditional centromere construct *P*_*GAL1*_*-CEN7* was used to replace the *CEN7* sequence on chromosome VII for transiently blocking disjunction of chromosome VII in the presence of galactose. In the *ura3Δ3’-HIS3-ura3Δ5’* cassette, a 390 bp direct repeat of a portion of *ura3* was designed in both *ura3Δ5’* and *ura3Δ3’* sequences. The galactose induction activates the *GAL1* promoter and blocks the function of *CEN7*, then the homologous recombination of the repeat of *ura3* can lead to excision and loss of *HIS3* and regeneration of functional *URA3*, which can lead to selection of aneuploid chromosome VII yeast cells YSy140 containing two copies of chromosome VII. *SynVII* was modified to carry the *LEU2* marker near the centromere, then through mating to YSy140 strain and sporulation, selection of His^+^Leu^+^ cells can lead to the final construction of aneuploid *synVII*. Whole genome sequencing of YSy142 revealed an extra copy of each chromosome I and III. By applying the same approach to generate YSy140, we successfully removed the extra copies of undesired chromosomes sequentially and leading to the auxotrophic marker on *chrVII* switched from *HIS3* to *MET15*. The chromosome sequence integrity of both *synVII* and chromosome VII was confirmed by whole genome sequencing on BGISEQ platform.

### Genome stability analysis of aneuploid *synVII* strain

The aneuploid *synVII* strains (strain ID: YSy140 and YSy142) were streaked on SC– Leu–Ura and SC–Leu–Met plates respectively and incubated at 30 °C for 5 days. Three single colonies of each strain were selected for successive subculture in 1mL SC medium for 210 generations at 30 °C, followed by plating on the SC plates and then replica plating was performed on both SC–Leu–Ura and SC–Leu–Met plates. The loss of synthetic or wildtype chromosome VII was calculated by counting numbers of colonies present on the SC plate but not on corresponding selective plates.

### SCRaMbLE and screening of aneuploid *synVII* strain

Aneuploid strains (strain ID: YSy142) with pSCW11*-CreEBD-URA3* plasmid were cultured in SC–Ura–Leu–Met liquid medium overnight at 30 °C. Cells were inoculated into SC–Ura–Leu–Met liquid medium with 1 μM β-estradiol and cultured for 24 hours at 30 °C. Then SCRaMbLEd cells were plated on SC–Leu–Met+0.01 μg/mL cycloheximide plates followed by 4 days incubation at 30 °C, with unSCRaMbLEd cells exposed in the same condition as phenotypic control. Single colonies with recovered phenotype compared to unSCRaMbLEd cells were selected for further phenotype tests and functional analyses.

### Chromosome copy number determination by quantitative real-time PCR (qPCR)

To determine the copy number of *chrI* and *chrIII*, an essential gene *YBR136W* on *chrII* was chosen as internal control, and *YAR007C, YCR052W* as targets on *chrI, chrIII* respectively. qPCR was conducted with 10ng genomic DNA in duplicate in a 20 μl reaction using the TB Green Premix Ex Taq II (Tli RNase H Plus, Takara) and the StepOnePlus™ Real-Time PCR System (ABI). Primer sequences: *YBR136W* forward primer 5’-TGGAACGTATTGGGGCTGAC-3’, reverse primer 5’-AGTCAGCGTCTG CTTGTTCA-3’; *YAR007C* forward primer 5’-GTGTGACGGATTTTGGTGGC-3’, reverse primer 5’-TGATGAAGTTTGCGTTGCGG-3’; *YCR052W* forward primer 5’-TTCCAACGATGCCGAAGACA-3’, reverse primer 5’-ACTTCACCCAATTCGG GCTT-3’.

### Zymolyase sensitivity analysis assay

Yeast cells were pre-cultured for 24 hours in minimal selective medium, and collected by centrifugation (3000 *g*, followed by Zymolyase treatment (20 μg/mL) either in 10mM Tris-HCl (pH 7.5) or in 0.8 M sorbitol (hyperosmotic). Cells were plated in 100-well honeycomb plate (catalog number: NC9976780) at 30 °C, and changes in optical density (OD) were measured at OD_600_ every 5 minutes with automatic Bioscreen C instrument (Helsinki, Finland) (n=3).

### Hi-C library construction

Hi-C libraries were generated using a protocol adapted from previous studies (Dekker et al., 2002; Mercy et al., 2017). 1 × 10^9^ cells were collected and washed from overnight cell culture, cells were cross-linked with 6 mL 3% formaldehyde in 1xPBS (137 mM NaCl, 2.7 mM KCl, 10 mM Na_2_HPO_4_, 1.8 mM KH_2_PO_4_) for 30 min, and quenched by adding 0.66 mL 2.5 M glycine (working concentration is 250 mM) at room temperature for 5 min and then keep on ice for 15 min. Cells were washed with 10 mL 1xPBS for 3 times and, pelleted cells were dissolved in 10 ml 1 M sorbitol with 5 mM DTT and 1 mg/mL Zymolyase 100T and incubated at 30 °C for 30-60 min to remove cell wall. Cells were lysed with 3 mL 0.2% Igepal CA-630 and 10 mM Tris-HCl, followed by 500 µL 0.5% SDS treatment at 62 °C for 10 min and 250 µL 10% TritonX-100 treatment at 37 °C for 15 min, then cells were washed and resuspended with restriction buffer (NEB3.1). Cross-linked DNA was digested at 37 °C overnight with 200 units of DpnII restriction enzyme (NEB) in a 500 µL reaction. The digestion mix was placed at 62 °C for 20 min, and put on ice immediately. DNA ends were repaired with 25 µL 0.4 mM biotin-14-dATP at 23 °C for 4 h, and ligated with 1000U NEB T4 ligase at 16 °C for 6 hours. DNA decrosslinking was performed through an overnight incubation at 65 °C with 25 µl 20 mg/ml proteinase K added to a total reaction volume of 1.2 mL. DNA was purified on XP beads, and unligated biotin ends were removed using T4 DNA polymerase. DNA was sheared and Biotin-tagged DNA were pull-down with Dynabeads MyOne Streptavidin T1 beads, repaired ends with T4 DNA polymerase and tagging A tails with NEB klenow exo minus, filled in adaptors with barcode oligo mix, PCR amplified resulting DNA and collected DNA with 300-600 bp size fragments, denatured DNA at 95 °C for 3 min and ssDNA cyclization with splint Oligo and T4 DNA ligase. The resulting Hi-C libraries were processed into whole genome sequencing on BGISEQ platform.

### Gene expression profiling analysis of aneuploid SCRaMbLEd synVII strains

The genome reconstruction of aneuploid SCRaMbLEd synVII was performed using the previously described method (Shen et al., 2016). The reference sequence of YSy142 was updated by adding the *synVII* sequence to the other chromosomes of the BY4741 reference genome.

For the transcriptome analysis, the environmental stress response (ESR) gene sets identified in previous studies (Brion et al., 2016; Gasch et al., 2000) were used for expression level analysis. The significance of GO terms in differentially expressed ESR genes was individually identified using the hyper-geometric test with false discovery rate (FDR) correction and the threshold P value < 0.001.

### Flow cytometry analysis

Asynchronous log-phase cells were fixed with 70% ethanol for 1 hour at room temperature. Then cell pellets were resuspended in 50mM sodium citrate (pH 7.0). Samples were briefly sonicated and placed on ice, followed by Rnase A (0.25 mg/mL) treatment for 1 hour at 50 °C. Cells were washed with 50 mM sodium citrate (pH 7.0) and resuspended into the same solution. Propidium iodide (16 µg/mL) was added to the cells and incubated at room temperature for 30 minutes. Samples were analyzed with BD FACSCelesta Cell Analyzer.

---

**Figure S1.**
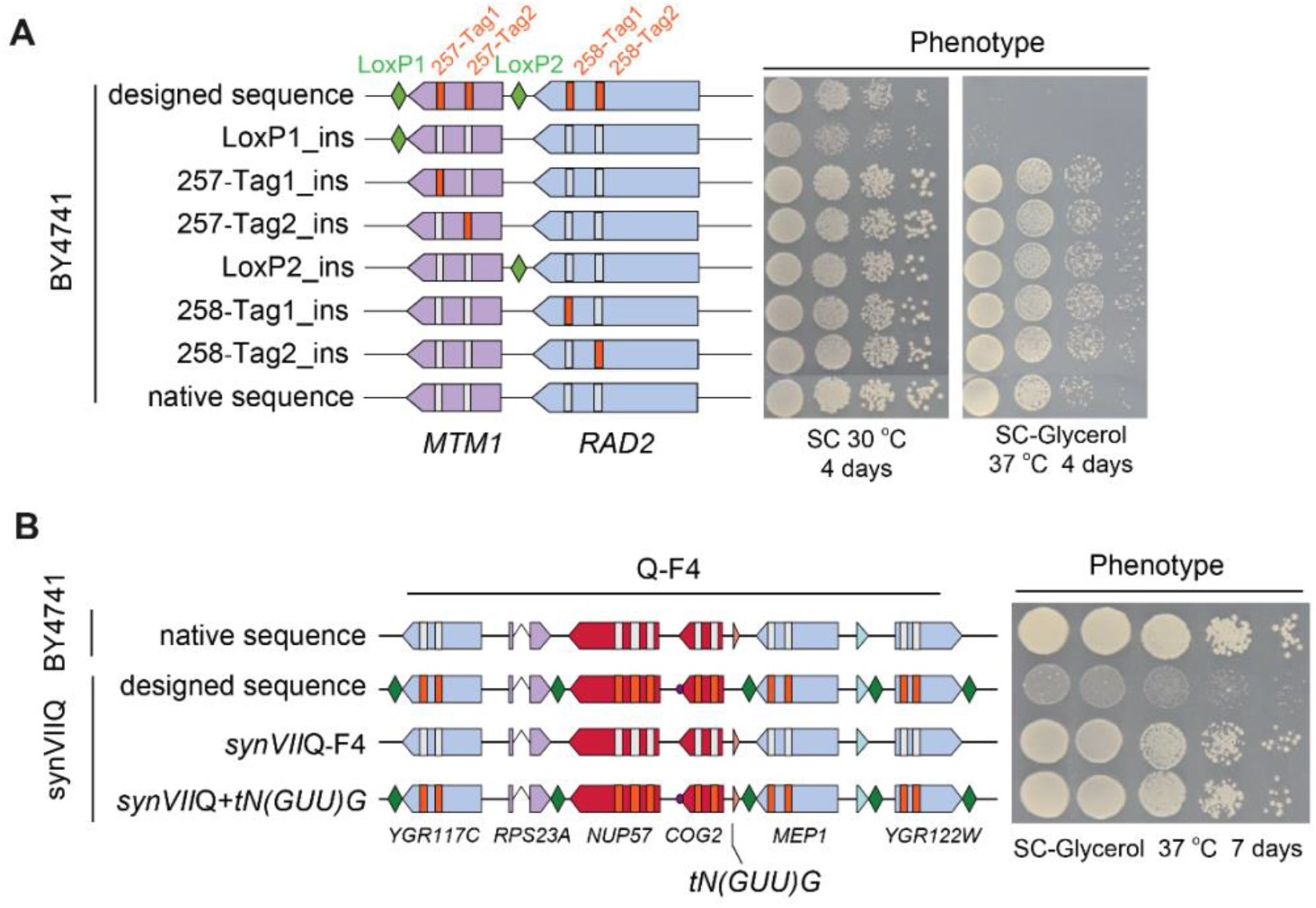
Debugging of W and Q regions in *synVII*. (A) Systematic dissection of defect origin in megachunk W by introducing each individual design feature into the BY4741 strain for phenotypic assay showing the loxPsym site at the 3’UTR of *MTM1* causes the defect. (B) Spotting assay with ten-fold serial dilutions of BY4741, synVIIQ intermediate strain and its derived strains with chunk 4 and tRNA gene *tN(GUU)G* in chunk 4 replaced by native sequence showing the addition of tRNA gene *tN(GUU)G* recovered the phenotype. The arrows represent gene order and orientations (red indicates essential genes, purple is fast-growth and blue represents non-essential). Green diamonds represent loxPsym sites embedded downstream of the stop codons. Vertical orange and white bars represent synthetic and wild-type PCRTags respectively.

**Figure S2.**
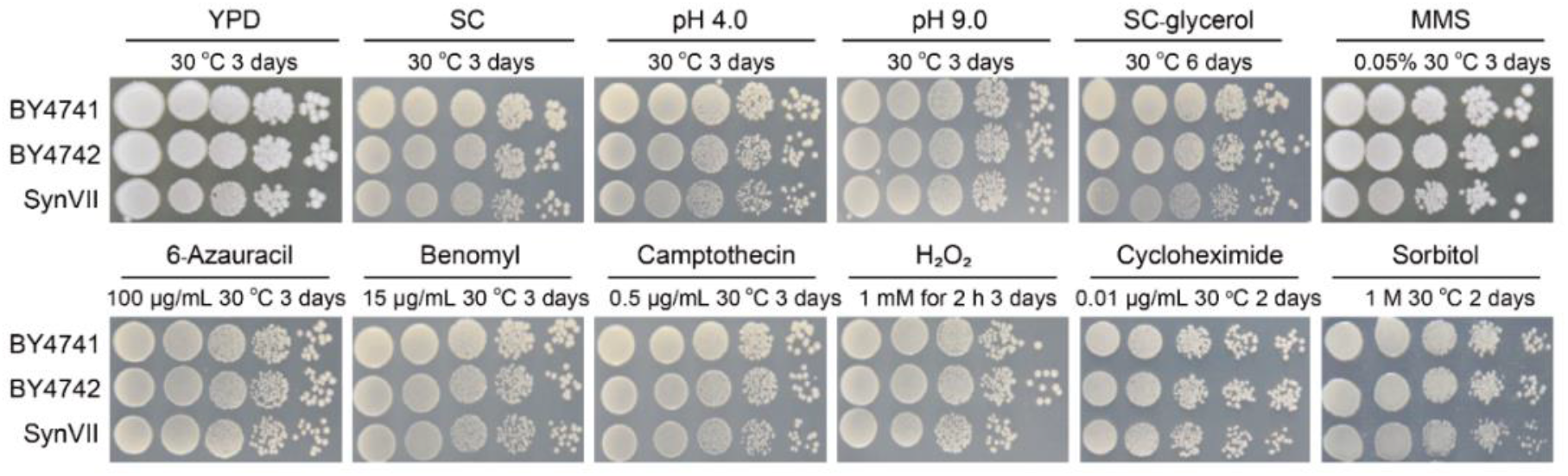
Phenotypic assays of synVII on different media in comparison with wild-type BY4741 and BY4742. 10-fold serial dilutions of overnight cultures of selected strains were used for plating. From left to right: YPD at 30 °C; SC at 30 °C; low pH YPD (pH 4.0); high pH YPD (pH 9.0); SC-Glycerol; YPD + MMS; SC + 6-azauracil; SC + benomyl; SC + camptothecin; SC + H_2_O_2_ (1 mM, 2 hours pretreatment); SC + cycloheximide; SC + sorbitol; (YPD, yeast extract peptone dextrose; SC, synthetic complete; MMS, methyl methane sulfone).

**Figure S3.**
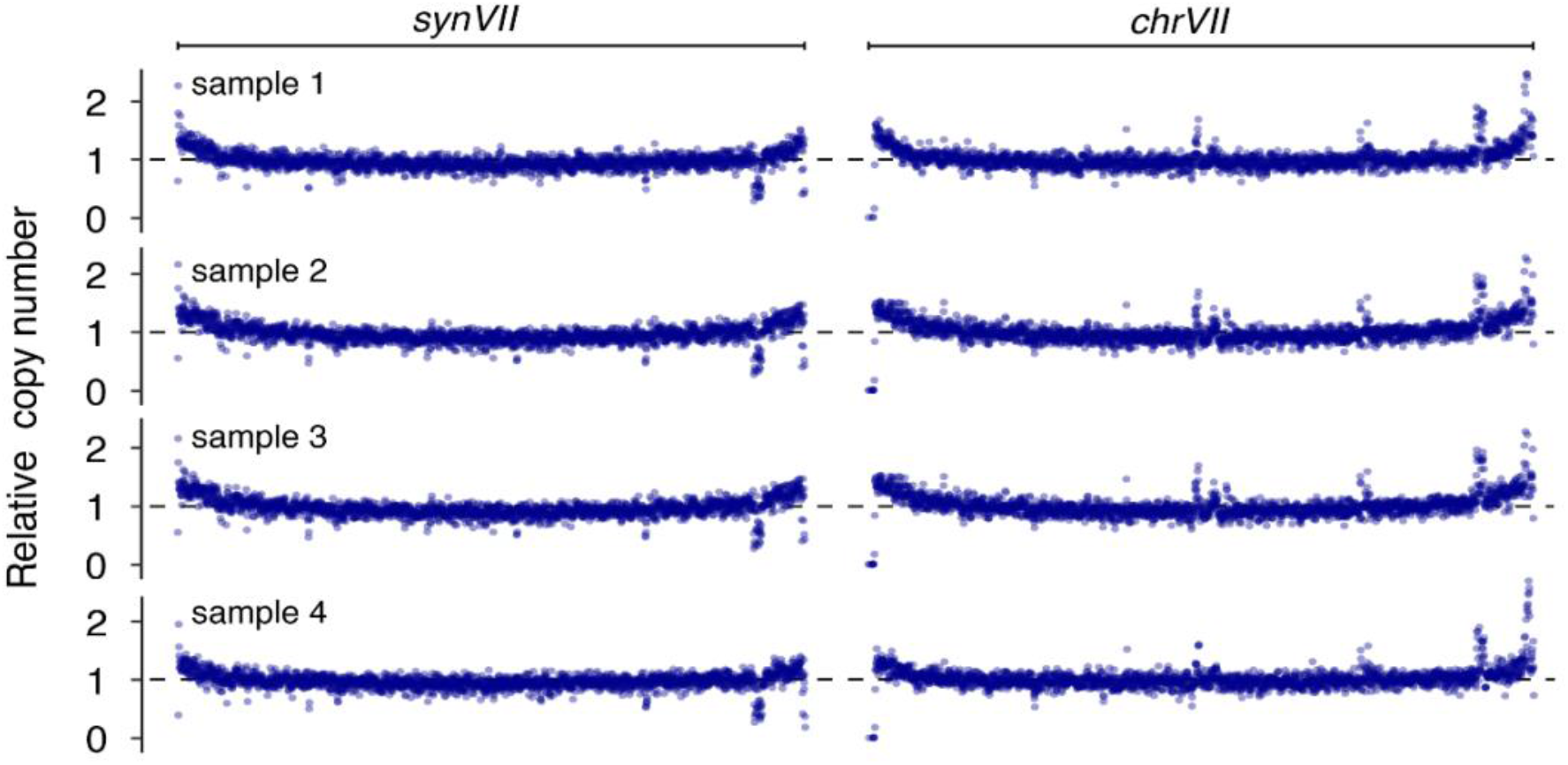
Verification of sequence integrity in disomic strain YSy142. The relative sequencing depth of YSy142 colonies after batch transfer for 220 generations. The relative sequencing depth was calculated in 500-bp windows by comparing to the average of sequencing depth of the whole genome.

**Figure S4.**
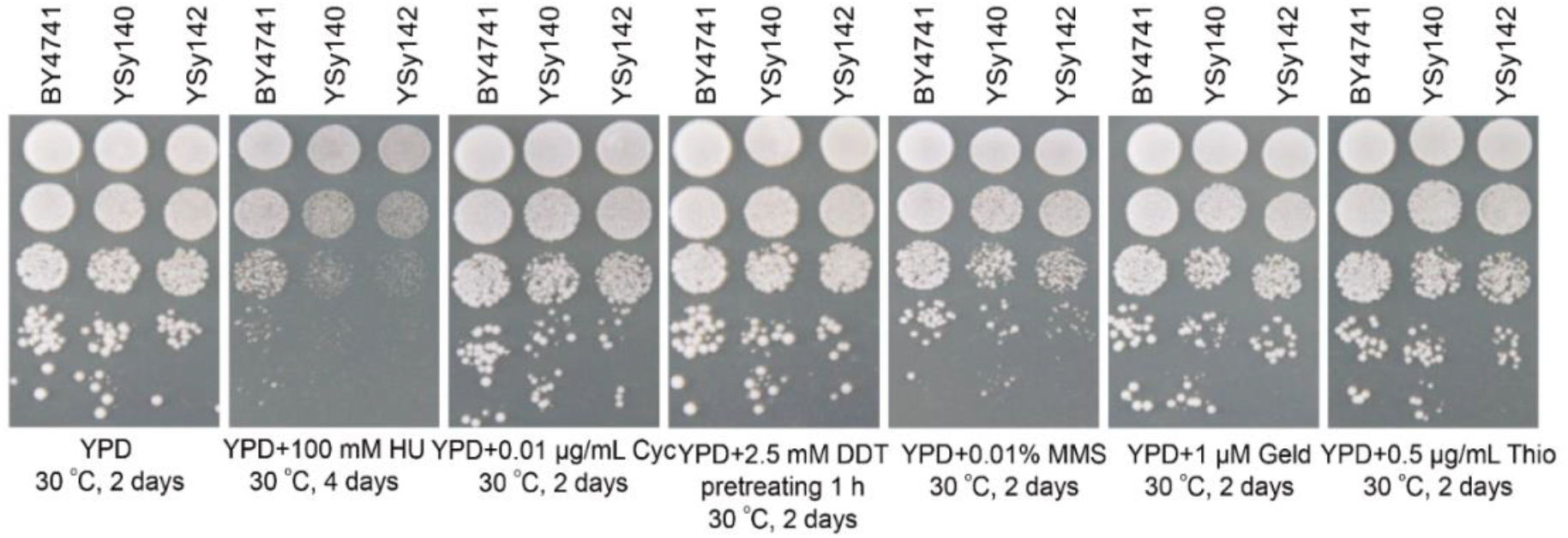
Spotting assay with ten-fold serial dilutions of disomic yeasts YSy140 and YSy142 compared to euploid strain BY4741 under selected conditions. From left to right: YPD at 30 °C; YPD + hydroxyurea (HU, 100 mM) at 30 °C; YPD + cycloheximide (Cyc, 0.01 μg/mL) at 30°C; YPD + DL-Dithiothreitol (DDT, 2.5 mM pretreating 1 h); YPD + methyl methane sulfone (MMS, 0.01% v/v) at 30°C; YPD + geldanamycin (Geld, 1 μM) at 30 °C; YPD + thiolutin (Thio, 0.5 μg/mL) at 30°C; (YPD, yeast extract peptone dextrose).

**Figure S5.**
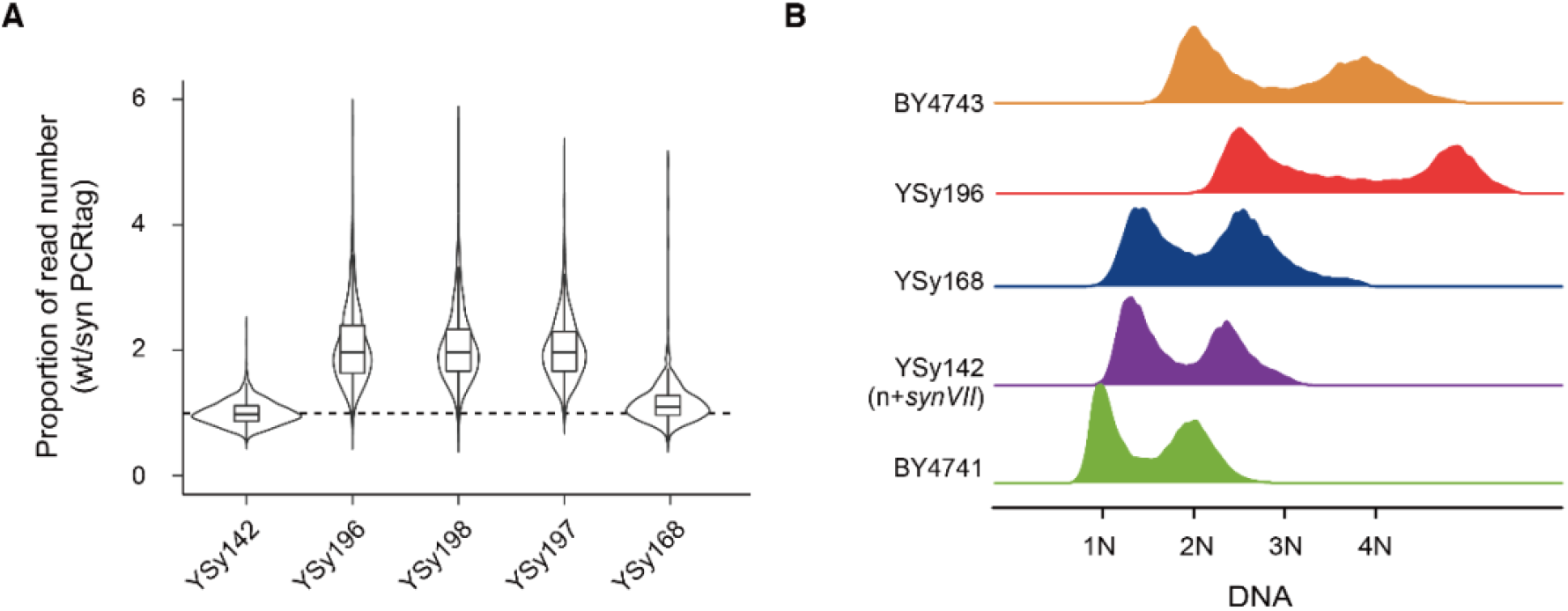
Karyotype analysis of SCRaMbLEd yeasts by whole genome sequencing analysis and flow cytometry. (A) The boxplot and violin plot present the proportion of reads for each wild-type (wt) versus synthetic (syn) PCRTags. The proportion is normalized by the mean of YSy142. The PCRtag sites of deleted region by SCRaMbLE is not included in the analysis. (B) For flow cytometry, asynchronous log-phase cells were analyzed (∼15000 to 18000 cells/each sample).

**Figure S6.**
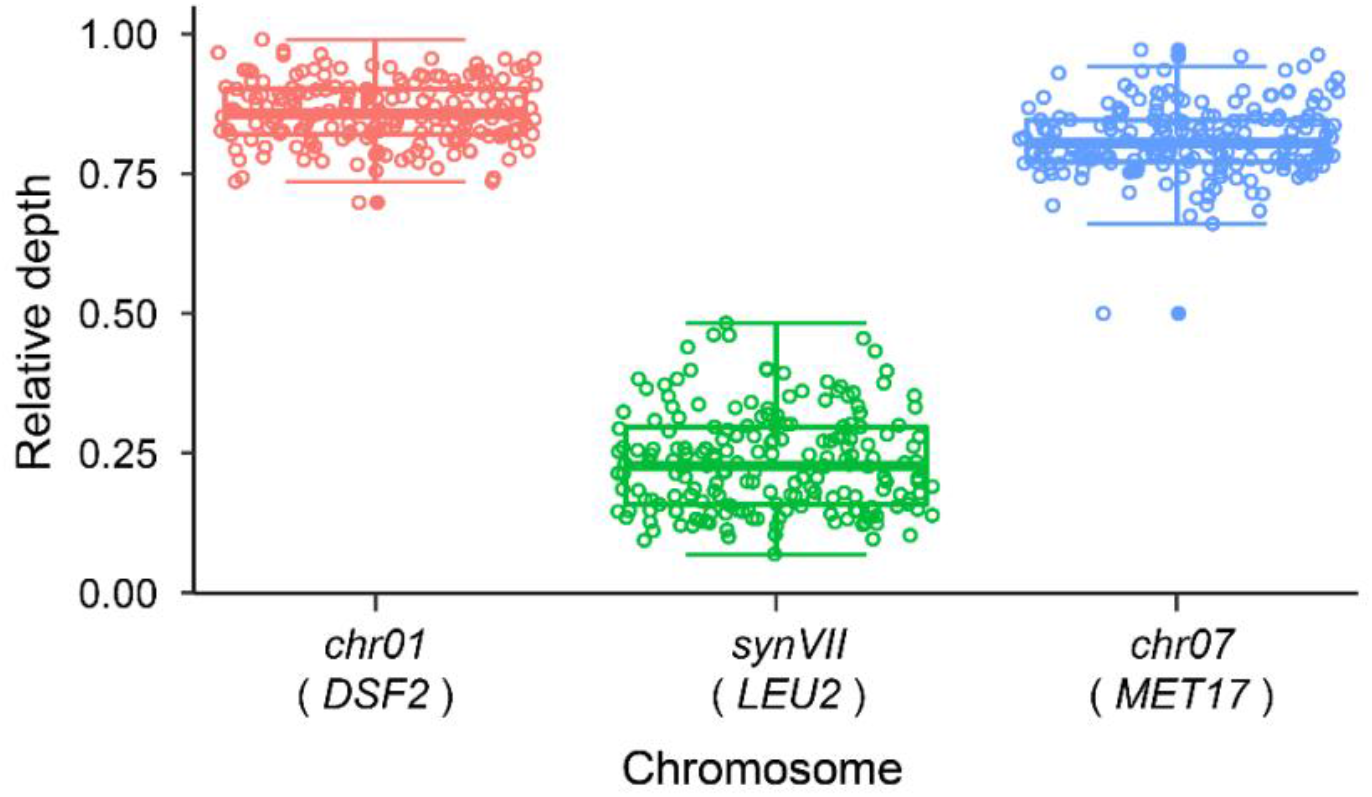
Relative average sequencing depth of 195 SCRaMbLEd strains with circular *synVII*. Deep sequencing coverage analysis revealed the lower depth of circular *synVII* in comparison with *chrVII* and *chrI* (control). Each dot represents one SCRaMbLEd strain. The relative depths were quantified by *DSF2* gene on *chrI, LEU2* gene on *synVII* and *MET17* gene on *chrVII* respectively. Each dot represents one SCRaMbLEd strain.

**Figure S7.**
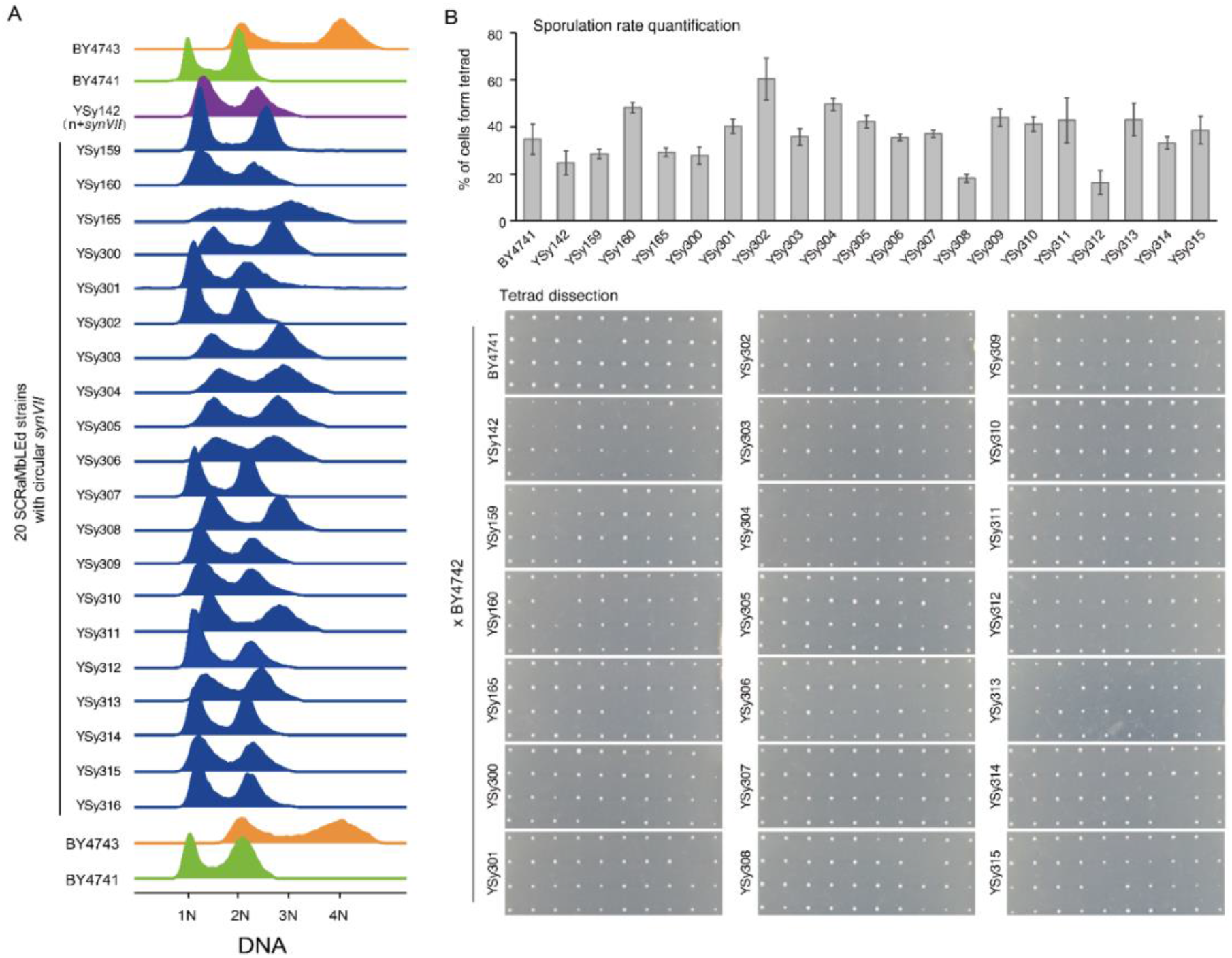
Karyotype validation by flow cytometry and sporulation in SCRaMbLEd strains with circular *synVII* chromosome. (A) For flow cytometry, ∼15000 to 18000 cells were sorted. (B) For sporulation all selected samples were first mated to BY4742, then followed by sporulation and tetrad dissection (for each sample, 300 cells were counted at least for sporulation rate quantification, 10 tetrads were selected for dissection).

**Figure S8.**
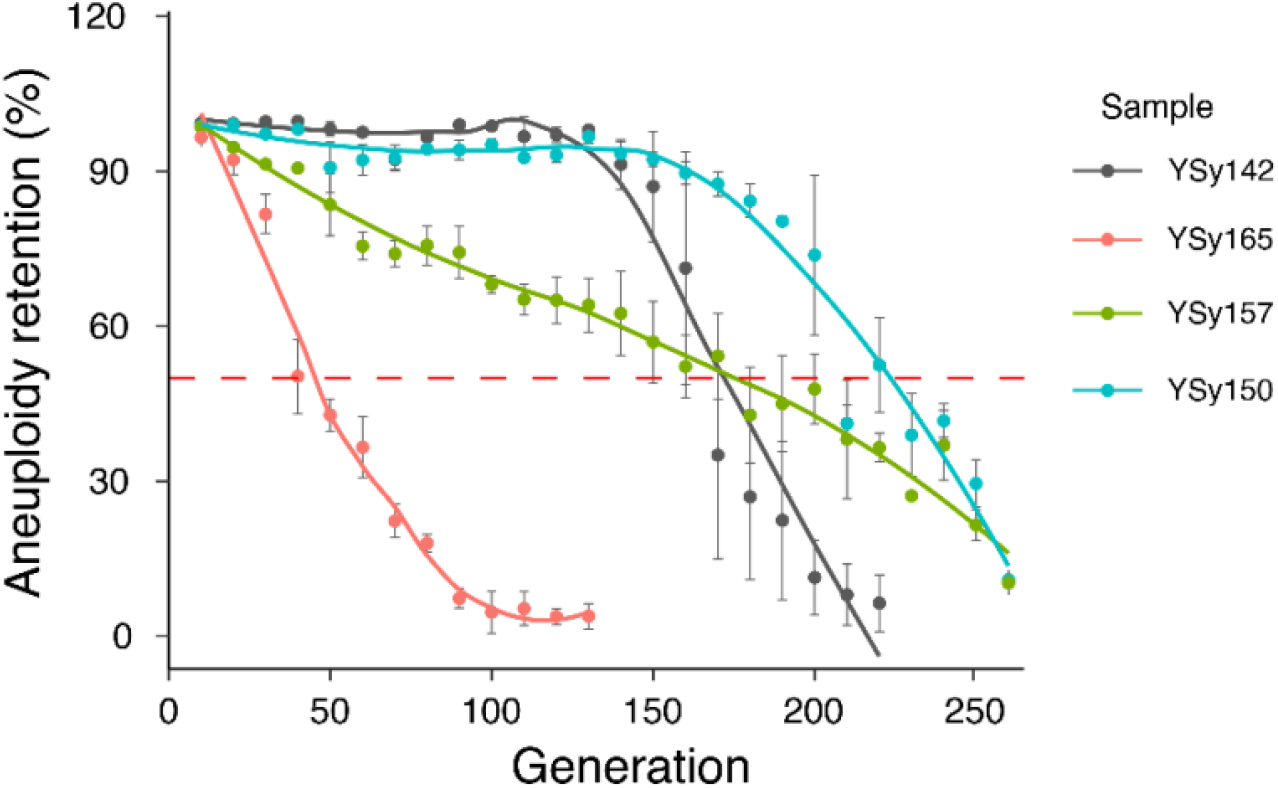
Genome stability analysis of SCRaMbLEd strains with circular *synVII* chromosome. Three representative strains with distinct *synVII* chromosome content and size were selected for genome stability assay by batch transfer for ∼230 generations. YSy142 (n + *synVII*) is used as control.

**Figure S9.**
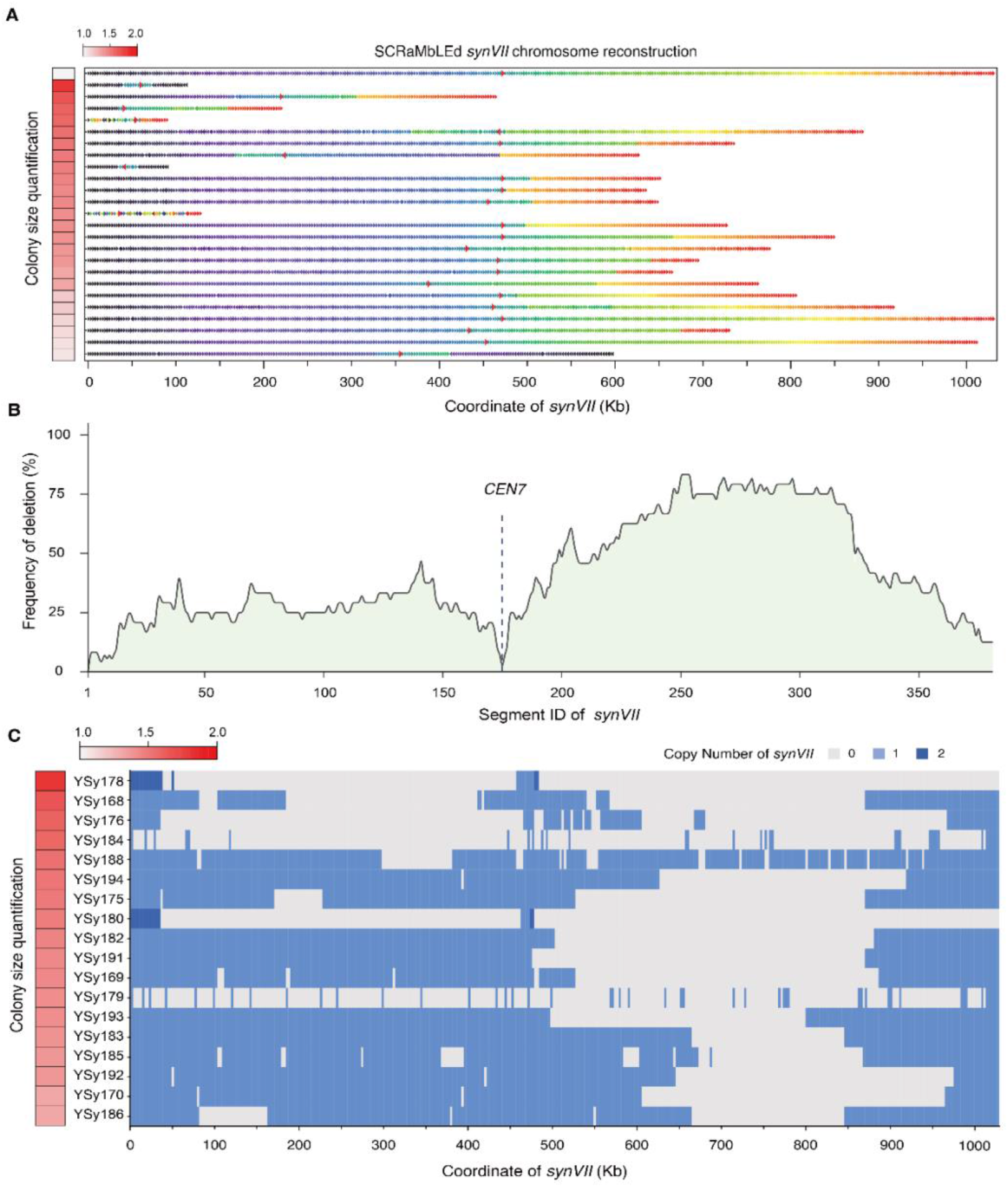
Deletion distribution across entire synthetic chromosome VII. (**A**) Rearrangements were observed in the 14 SCRaMbLEd aneuploid yeasts with linear *synVII*. Each SCRaMbLE strain is represented as a sequence of arrows (SCRaMbLE-gram). The color and direction of each arrow indicates the segment number in the parental chromosome and its orientation. A red border denotes a segment containing *CEN7*. Y-axis shows the relative average phenotypic recovery rate of each SCRaMbLEd strain in comparison to that of YSy142 represented by color scale (n ≥ 200). (**B**) Deletion hotspot observed in the right arm region of *synVII*. Y-axis represents the percentage of SCRaMbLEd strains that with corresponding segment been deleted. (**C**) The fate of each segment flanked by two loxPsym sites in each strain is indicated as deleted (gray) and one copy with no change (light blue).

**Table S1.** Summary of *synVII* design.

| Modification | Design feature | Number | Base alteration |
| --- | --- | --- | --- |
| Replacement | wild type telomere -> Universal Telomere Cap<sup>a</sup> | 2 | 21094 -> 1378 |
|  | 705 pairs of WT PCR tags and 705 pairs of SYN PCRTags | 1410 | 17876 |
|  | stop codon TAG -> TAA | 126 | 126 |
|  | “landmark” restriction sites (removal or introduction)<sup>b</sup> | 261 | 596 |
|  | repeatsmash of gene | 3 | 2147 |
| Deletion | transposable element region<sup>c</sup> | 20 | 51111 |
|  | gene | 14 | 12232 |
|  | tRNA | 36 | 2887 |
|  | intron<sup>d</sup> | 23 | 3062 |
| Insertion | LoxPsym site | 380 | 12920 |

a. Wild-type telomeres together with its adjacent subtelomeric regions are replaced with universal telomere caps.
b. Unique restriction sites were generated in *synVII* either by introduction of synonymous mutations in ORFs, or by removal of redundant sites by the same method to leave only one pre-existing instance of a given site in the *synVII* sequence.
c. Transposable element region include these functional features of LTR retrotransposons, transposable element genes and long terminal repeats.
d. Eleven introns were retained as these introns reside in ribosomal subunit coding genes.

**Table S2.**
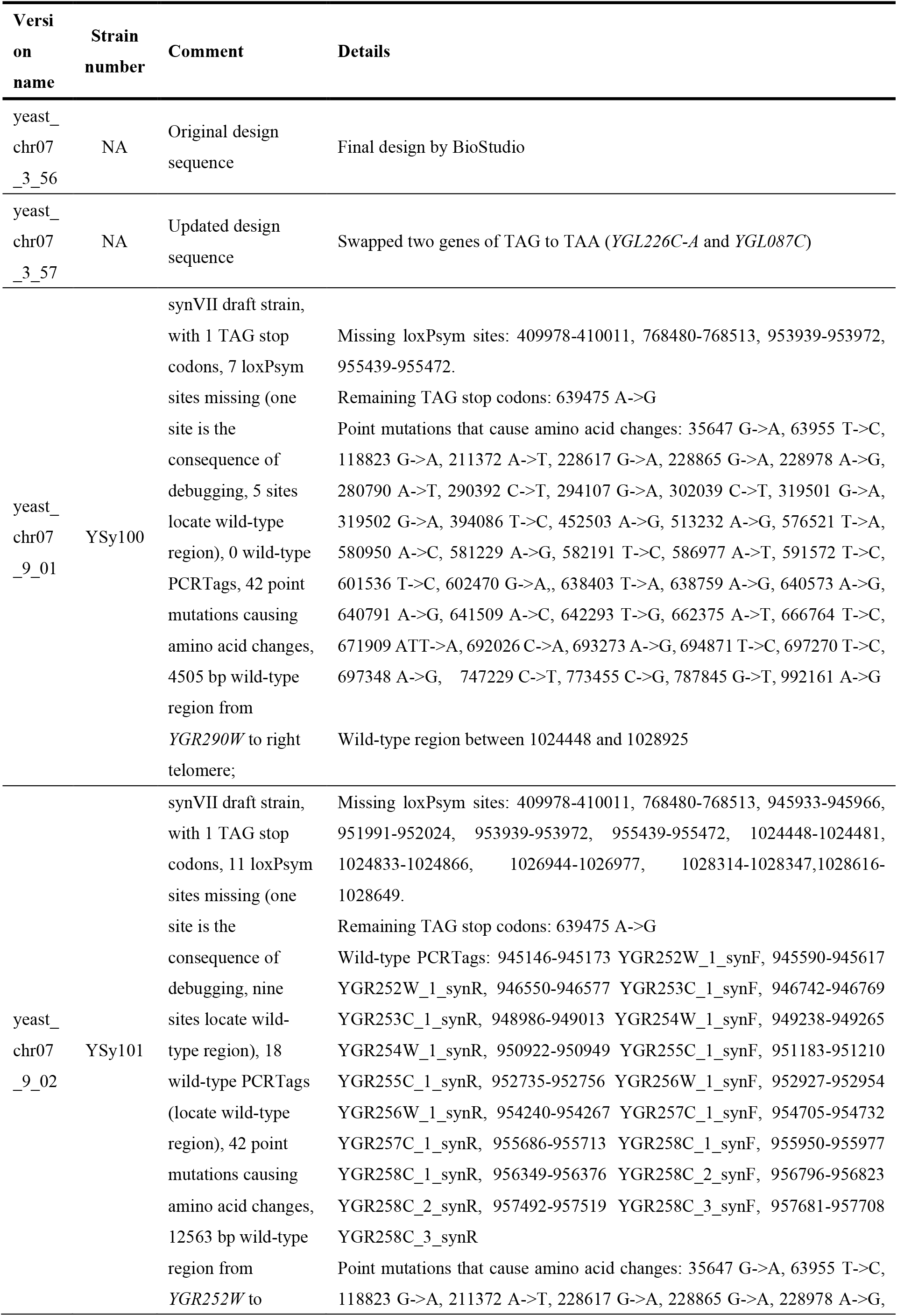
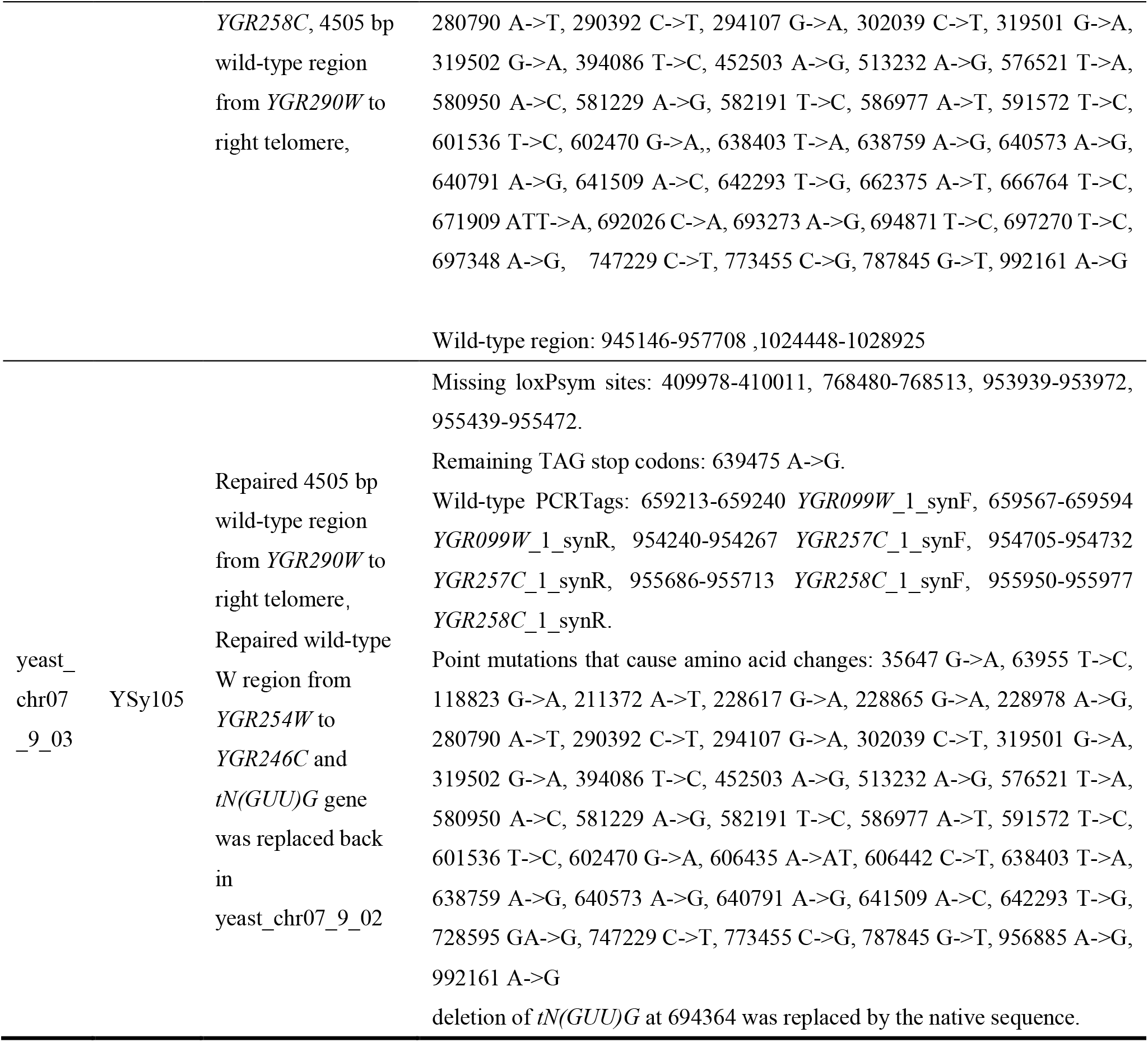
Chromosome version used in this paper.

**Table S3. Yeast strains used in this paper**

